# The mitochondrial chaperone HSPD1 folds MTHFD2 independently of its co-chaperone HSPE1

**DOI:** 10.64898/2026.02.16.706072

**Authors:** Shani Gabbay, Hila Ben-David, Shatha S Alassam, Linor Cohen, Tal Levy, Liron Levin, Nili Tickotsky Moskovitz, Ifat Abramovich, Albert Batushansky, Shiran Dror, Moshe Elkabets, Noga Alon, Yariv Brotman, Shai Kaluski-Kopatch, Evelina Nikelshparg, Sufa Sued-Hendrickson, Shimon Bershtein, Anat Ben-Zvi, Barak Rotblat

**Affiliations:** Department of Life Sciences, Ben-Gurion University of the Negev, Beer-Sheva, Israel; Ilse Katz Institute (IKI) for Nanoscale Science and Technology, Ben-Gurion University of the Negev, Beer Sheva, Israel; Laura and Isaac Perlmutter Metabolomics Center, B. Rappaport Faculty of Medicine, Technion-Israel Institute of Technology, Haifa, Israel; The Shraga Segal Department of Microbiology, Immunology and Genetics, Faculty of Health Science, Ben-Gurion University of the Negev, Beer-Sheva, 84105, Israel; Faculty of Health Sciences, Ben-Gurion University of the Negev, Beer-Sheva, 84105, Israel

**Author notes:** Equal contribution (Shani Gabbay, Hila Ben-David).

## Abstract

Acquiring new cellular states entails metabolic reprogramming driven by changes in the expression of cytosolic and mitochondrial metabolic enzymes. Most mitochondrial proteins are synthesized in the cytosol and imported into the mitochondria in a linear form, after which they are folded by a network of mitochondrial chaperones and co-chaperones. Which mitochondrial protein is dependent upon which chaperone for its folding is largely unknown. HSPD1/HSPE1 (HSP60/HSP10) are evolutionarily conserved mammalian homologues of the bacterial proteins GroEL/GroES, forming a chamber-and-lid chaperonin to facilitate the folding of client proteins. We used gene knockdown and SILAC-based proteomics to identify HSPD1 client proteins. We found that HSPD1 supports the expression of Methylenetetrahydrofolate Dehydrogenase 2 (MTHFD2), a key essential protein in the mitochondrial one-carbon (1C) pathway, in cells and tumors, and directly folds MTHFD2, independently of its co-chaperone HSPE1. HSPD1 interacts with MTHFD2 in mitochondria, and MTHFD2 is degraded by LONP1 in HSPD1 knockdown cells. Consequently, we observed reduced nucleotide and S-adenosylmethionine (SAM) levels in HSPD1 knockdown and found minimal overlap in the transcriptional and metabolic cellular responses to HSPD1 vs. HSPE1 depletion. In C. elegans, knockout of HSP60 triggers the mitochondrial stress response in the gut, while HSP10 knockout triggers the mitochondrial stress response in muscle tissue. Our data support that HSPD1 is an MTHFD2 chaperone and that, in addition to working together, HSPD1 and HSPE1 have distinct biological functions.

## Introduction

Metabolic reprogramming is the process by which a cell changes its metabolic profile, such as during differentiation^1^, immunity^1,2^, and response to environmental cues^3^. In addition, metabolic reprogramming is a hallmark of cancer and a crucial aspect of the process by which a normal cell becomes malignant and metastatic^4–7^. Metabolic reprogramming entails the up- and down-regulation of enzymes in the cytosol and mitochondria^8,9^. Most mitochondrial enzymes are encoded in the nuclear genome and synthesized in the cytosol. These enzymes are imported into the mitochondria in an unfolded state and must be correctly folded by mitochondrial chaperones^10^. Mitochondrial chaperones operate within sophisticated networks alongside other chaperones and co-chaperones to facilitate precise protein folding. Additionally, they fold stress-denatured mitochondrial proteins. When the stoichiometric balance between chaperones and their client proteins is disrupted, the consequences include protein misfolding, aggregation, or premature degradation^11^.

There is accumulating evidence that specific proteins depend on specific chaperones for folding. For example, the mitochondrial chaperone, TRAP1, stabilizes the electron transport chain Complex II subunit succinate dehydrogenase-B, along with other mitochondrial proteins, to facilitate respiration during nutrient stress, supporting the possibility that specific mitochondrial chaperones are critical for specific mitochondrial functions^12^. In line with this concept, the expression of specific chaperones and their clients is coordinated in tissues during development, exemplified by up-regulation of the myosin chaperon Unc45B during muscle development in *C. elegans*, indicating that molecular mechanisms have evolved to ensure sufficient chaperone capacity to accommodate the folding of proteins expressed in the particular biological context^13^. Using gene expression data obtained from patient-derived tumors, we previously found that mitochondrial chaperones and proteins form a network comprised of two major clusters^14,15^, suggesting that coordinated expression of proteins with their chaperones evolved to provide sufficient chaperone expression to support the folding of these proteins.

HSPD1 and HSPE1 (HSP60 and HSP10) are chaperone-co-chaperone complexes, homologs of bacterial GroEL-GroES, together forming a protein folding chaperonine^16^. HSPD1 and HSPE1 reside in the mitochondrial matrix, catering to the folding of newly imported and synthesized proteins as well as stress-induced misfolded proteins. The HSPD1 interactome was studied using crosslinking and immunoprecipitation, and more than 300 proteins were identified in complex with HSPD1, raising the question of which of these depend on HSPD1 for their folding in cells^17^. Treating cells with the small-molecule inhibitor of HSPD1, KHS101, led to the aggregation of mitochondrial proteins involved in glycolysis, the TCA cycle, OXPHOS, and mitochondrial quality control^18^. In addition, KHS101 reduced tumorigenicity in glioblastoma and lung cancer tumor cells in vitro and in vivo^19^. Furthermore, HSPD1 depletion led to reduced proliferation and oxygen consumption in lung cancer cells^19^. Conversely, HSPD1 knockdown (KD) in ovarian tumor cells resulted in increased proliferation by promoting the expression of the fatty acid elongation enzyme 3-Oxoacyl-ACP Synthase (OXSM)^20^, implicating HSPD1 in fat metabolism. Recently, HSPD1, together with its co-chaperone HSPE1, was found to play essential roles for the expression of mitochondrial ribosomal proteins and, therefore, for mitochondrial translation^21^. Nevertheless, it is not clear which of the HSPD1-dependent proteins directly interact with HSPD1 in the mitochondria or can be directly folded by HSPD1. Additionally, the metabolic pathways that depend on HSPD1 for their function are not fully understood.

While HSPE1 is assumed to be indispensable for HSPD1 function as a molecular chaperone^21,22,23,24^ KD of HSPE1, but not HSPD1, affects the processing of a mitochondrial GTPase, suggesting that, in addition to functioning as a complex together, the two proteins have distinct cellular functions^25,26^. Network analysis of mitochondrial chaperone-protein coexpression in cancer indicates that HSPD1 and HSPE1 are coexpressed with different sets of mitochondrial proteins^14,15^. In addition, the expression of each protein is correlated with sensitivity to drugs targeting distinct pathways in tumor cells^27^. These data support the premise that in mammalian cells, HSPE1 may be required for HSPD1-mediated folding of only a distinct set of its clients. The cellular functions of HSPD1 and those that are HSPE1-independent, in particular, are not well known.

Here, we found that HSPD1 folds MTHFD2 independently of HSPE1 and that HSPD1 KD leads to LONP1-dependent MTHFD2 degradation. Functionally, HSPD1 KD but not HSPE1 KD interacts with nucleotide and SAM metabolism, both of which are linked to the 1C pathway. In c. elegans, knockout of HSP60 triggers the mitochondrial unfolded protein response UPRmt in the gut, while knockout of HSP10 triggers UPRmt in muscles, supporting that each protein evolved a key function in a distinct biological context.

## Results

### HSPD1-dependent mitochondrial proteins

We hypothesized that a subset of mitochondrial proteins depends on HSPD1 for folding. We reasoned that HSPD1 depletion would lead to aggregation or degradation of such HSPD1-client proteins. To test our hypothesis and identify HSPD1-dependent proteins, we attempted to generate HSPD1 knockout (KO) U251 cells, in which HSPD1 was found to be functional^18^, using CRISPR. HSPD1 is an essential gene^28^, which may explain why we failed to obtain single-cell-derived clones with complete ablation of HSPD1. To generate cells where HSPD1 is depleted by >80%, we chose clones with reduced HSPD1 expression and used lentivirus to stably express shRNA targeting HSPD1 (two independent sequences). Cells expressing scrambled shRNA (shCONT) were our controls (Fig. 1a).

**Fig. 1.**
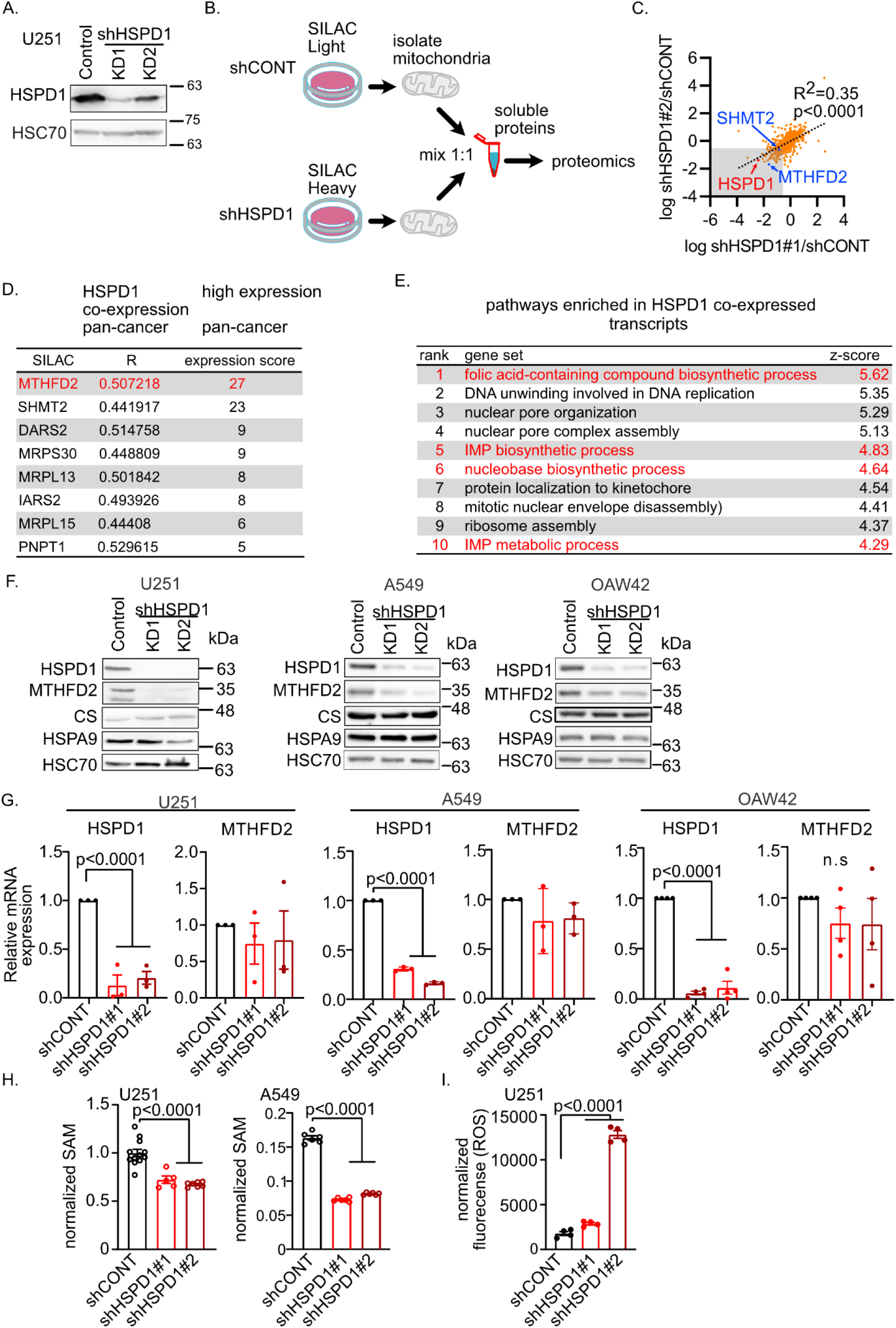
HSPD1 KD impacts MTHFD2. A. Lysates of HSPD1 KD and control cells were analysed using western blot. Ponceau staining was used as a loading control. B. Scheme depicting the SILAC experiment. C. SAILC ratio of mitochondrial proteins in each HSPD1 KD vs. control cells are plotted. The gray shaded area depicts proteins down-regulated by >50% in both KD cells vs. the control. D. proteins ranked according to their co-expression with HSPD1 and expression level in different cancer types. E. ARCHS4 analysis of enriched pathways in transcripts co-expressed with *HSPD1* across multiple RNAseq datasets. F. Lysates from HSPD1 KD and control cells were analysed by western blot. HSC70 was used as a loading control. G. qRT-PCR was used to measure the expression of the indicated transcripts in HSPD1 KD and control cells. H. SAM levels in HSPD1 KD and control cells. I. ROS levels were determined using DCFDA and FACS.

We took a proteomics approach to identify mitochondrial proteins that exhibit reduced solubility or expression in HSPD1 KD cells, as is expected in cases that depend on HSPD1 for their folding. We used SILAC and labeled proteins of HSPD1 KD cells with heavy lysine and arginine and proteins of control cells with normal amino acids. We isolated mitochondria from cell lysates using anti-TOM22 beads (MACS), lysed the mitochondria, and mixed the soluble fractions in a 1:1 ratio for proteomics analysis (Fig. 1b). We found 107 proteins whose expression was reduced in the two KD cell lines using -log2<0.5 as a cutoff (Fig. 1c; Table S1). We interrogated the SILAC list of down-regulated proteins using Enricher^29–31^ and found enrichment for tRNA Aminoacylation and One Carbon (1C) Metabolism pathway in Reactome Pathways and WikiPathways 2024 Human, respectively (Table S2).

Our model predicts that folding might be a limiting factor in cases where the chaperone’s client is highly expressed, burdening the client’s chaperones, and, therefore, the expression of the client and chaperone will be coordinated. To prioritize proteins for further analysis, we chose proteins whose expression is highly correlated with HSPD1 and is highly expressed in multiple cancer types. To rank proteins according to their expression levels in multiple cancer types and their co-expression with HSPD1, we utilized previously published analyses of metabolic enzyme expression in cancer^32^ and mitochondrial chaperone-protein co-expression in multiple cancer types^14^ (Fig. 1d). MTHFD2 was chosen for validation as it exhibited high co-expression with HSPD1 (ranked #3) and is highly expressed in multiple cancer types (#1). In addition, using the ARCHS4 database to interrogate more than 100,000 RNA-seq experiments^33^, we found that HSPD1 is mostly co-expressed with the 1C pathway proteins (Fig. 1e), supporting the possibility that HSPD1 promotes the pathway.

MTHFD2 is a key mitochondrial enzyme in the 1C pathway. The mitochondrial 1C pathway provides carbons for nucleotide synthesis^34,35^, SAM generation^36^, and reducing power for antioxidants (NADPH)34, making it a promising drug target in oncology^37^. We identified another key mitochondrial 1C enzyme, SHMT2, as downregulated in our SILAC experiment (Fig. 1c). SHMT2 and MTHFD2 work in concert within the mitochondrial 1C pathway. SHMT2 generates 5,10-CH₂-THF, a substrate for MTHFD2. SHMT2 is co-expressed with HSPD1 (ranked #6) and is highly expressed in cancer (#2) (Fig. 1d).

Using western blot, we confirmed reduced expression of MTHFD2 in HSPD1 KD cells and sustained expression of CS and HSPA9, mitochondrial proteins that did not change significantly in our SILAC experiment (Fig. 1f). We validated MTHFD2 downregulation, but not CS or HSPA9, upon HSPD1 KD in two other cell lines (Fig. 1g) in which HSPD1 was found to be functional, A549^19^ and OAW42^20^.

The reduced expression of MTHFD2 protein in HSPD1 KD cells may result from decreased MTHFD2 mRNA levels in cells coping with reduced HSPD1 levels, rather than being due to MTHFD2 misfolding, aggregation, and/or degradation. To rule this possibility out, we used RT-qPCR to measure HSPD1 and MTHFD2 mRNA levels in HSPD1 KD and control cells. We found reduced *HSPD1* but not *MTHFD2* mRNA levels in HSPD1 KD cells (Fig. 1g) and concluded that reduced MTHFD2 protein found in HSPD1 KD cells is not due to reduced *MTHFD2* mRNA levels. These results suggest that MTHFD2 is a client protein of HSPD1 and that HSPD1 may play an important role in the 1C pathway.

The 1C pathway is essential for the generation of a key methyl donor, S-Adenosylmethionine (SAM)^38,39^. We therefore opted to measure SAM as a surrogate for the 1C output. We synthesized heavy SAM, spiked our lysis buffer with it, and used it as a standard to quantify SAM in HSPD1 KD and control cells. We found that, indeed, SAM levels are reduced in HSPD1 KD U251 cells and found that this is the case in A549 cells as well (Fig. 1H). The mitochondrial 1C in general and MTHFD2 in particular generate NADPH, a major cellular antioxidant, and MTHFD2 depletion leads to increased cellular ROS^40,41^. In agreement, we found increased ROS in HSPD1 KD cells (Fig. 1i). Together, these results support that HSPD1 interacts with the 1C pathway.

Since HSPD1 is an important chaperone and essential gene, our results using HSPD1 KD cells might reflect total mitochondrial collapse rather than misfolding of a specific subset of HSPD1 clients. To this end, we used Seahorse XF to measure mitochondrial respiration in HSPD1 KD and control cells (Fig. S1a and b). We did not find consistent differences in basal or maximal mitochondrial respiration following HSPD1 KD in U251 cells (Fig. S1a); however, HSPD1 KD did reduce basal and maximal mitochondrial respiration in A549 cells (Fig. S1b). In addition, we measured mitochondrial mass using mitotracker green and found that HSPD1 KD did not affect mitochondrial mass in U251 or A549 cells (Fig. S1C). Since mitochondrial mass and respiration were intact in HSPD1 KD cells and since HSPD1 depletion led to reduced expression of MTHFD2 but not CS or HSPA9, we can conclude that mitochondria did not collapse during the time of our experiments, supporting that our positive SILAC hits represent putative HSPD1 client proteins.

### MTHFD2 interacts with HSPD1 in mitochondria

We next hypothesize that HSPD1 folds MTHFD2 in the mitochondria. While co-IP experiments found that the two proteins interact in cell lysates^17^, it is unclear whether they interact in intact cells. We used Proximity Ligation Assay (PLA), which detects potential protein-protein interaction (<40nm proximity) in fixed cells using anti-HSPD1 and MTHFD2 antibodies. We found a clear perinuclear PLA signal in control cells, which was reduced in HSPD1 KD cells (Fig. 2a). Using MitoTracker RED to stain mitochondria, we found PLA signal localized in the mitochondria (Fig. 2b), supporting that HSPD1 and MTHFD2 interact, or are at least in close proximity to each other, in the mitochondria.

**Fig. 2.**
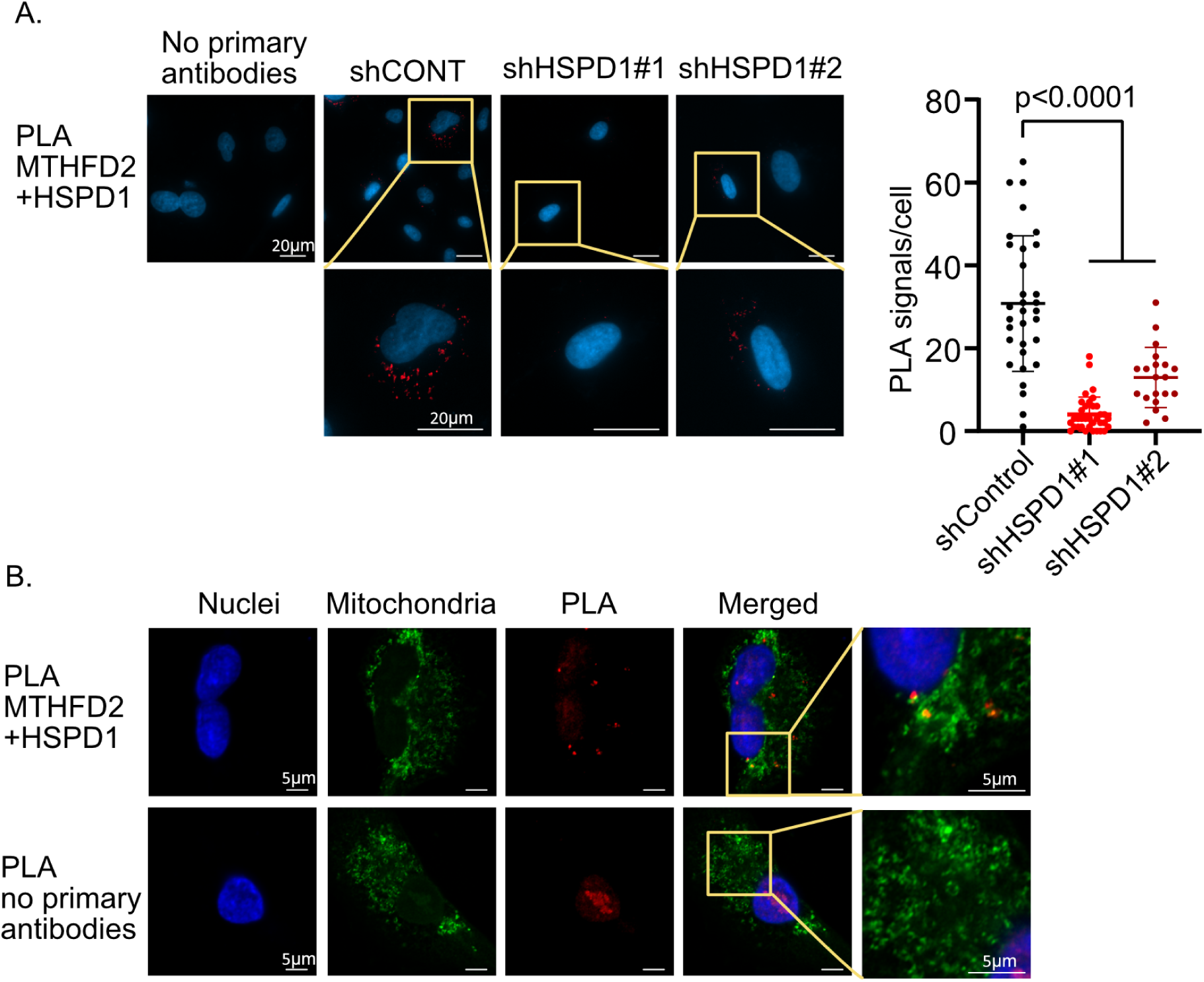
HSPD1 interacts with MTHD2 in mitochondria. A. PLA analysis of HSPD1 and MTHFD2 in A549 cells was performed using HSPD1 KD cells and no primary antibodies as controls. Dots were counted in at least 40 cells. B. PLA of HSPD1 interaction with MTHFD2 (red) and mitochondria staining (green).

### HSPD1 promotes MTHFD2 in tumors

We used a mouse xenograft model to test whether HSPD1 promotes MTHFD2 in tumors. HSPD1 KD and control A549 cells were injected into the flanks of NOD-SCID mice. Tumors were excised after thirty-eight days, and the levels of HSPD1 and MTHFD2 were determined using western blot (Fig. 3a; Sup Fig. 2a). We found reduced HSPD1 and MTHFD2 in tumors generated by HSPD1 KD cells (Fig. 3a, Sup Fig. 2a), indicating that HSPD1 promotes MTHFD2 in tumors.

**Fig. 3.**
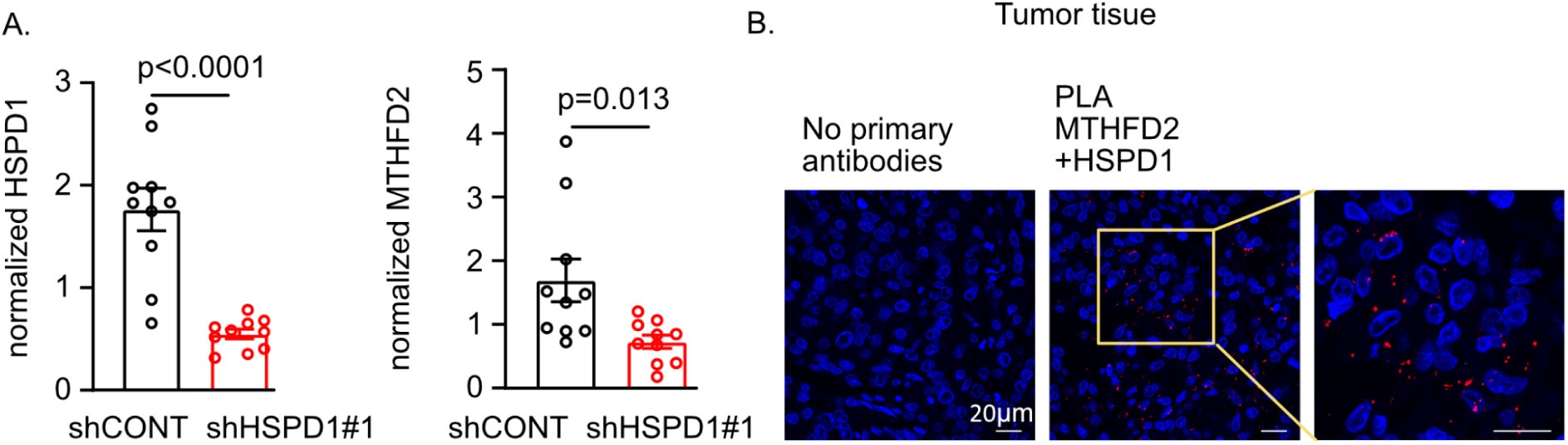
HSPD1 promotes MTHFD2 in tumors. A. HSPD1 and MTHFD2 levels in tumors generated by HSPD1 KD and control A549 cells were measured using western blot and normalized to loading control. B. PLA between HSPD1 and MTHFD2 in A549 tumors generated by shCONT cells.

To determine if HSPD1 and MTHFD2 interact in tumor tissue, we used PLA and shCONT tumor tissue. We found a clear PLA signal in the stained tissue (Fig. 3b), indicating that HSPD1 and MTHFD2 interact, or are at least in close proximity (<40 nm) to HSPD1 in tumor tissue. We conclude that HSPD1 interacts with MTHFD2 in tumor tissue to support its expression.

### MTHFD2 is degraded by LONP1 in HSPD1 KD cells

We asked if increased protein degradation rates can explain the reduced MTHFD2 levels found in HSPD1 KD cells. We pulsed cells with cyclohexamide (CHX) to stop new protein synthesis and measured the levels of remaining MTHFD2 at different time points following treatment (Fig. 4a). We calculated MTHFD2 half-life using densitometry measurements of band intensities and plotting the ln values over time. We found reduced MTHFD2 half-life in HSPD1 KD cells, indicating increased MTHFD2 degradation rates in HSPD1 KD cells.

**Fig. 4.**
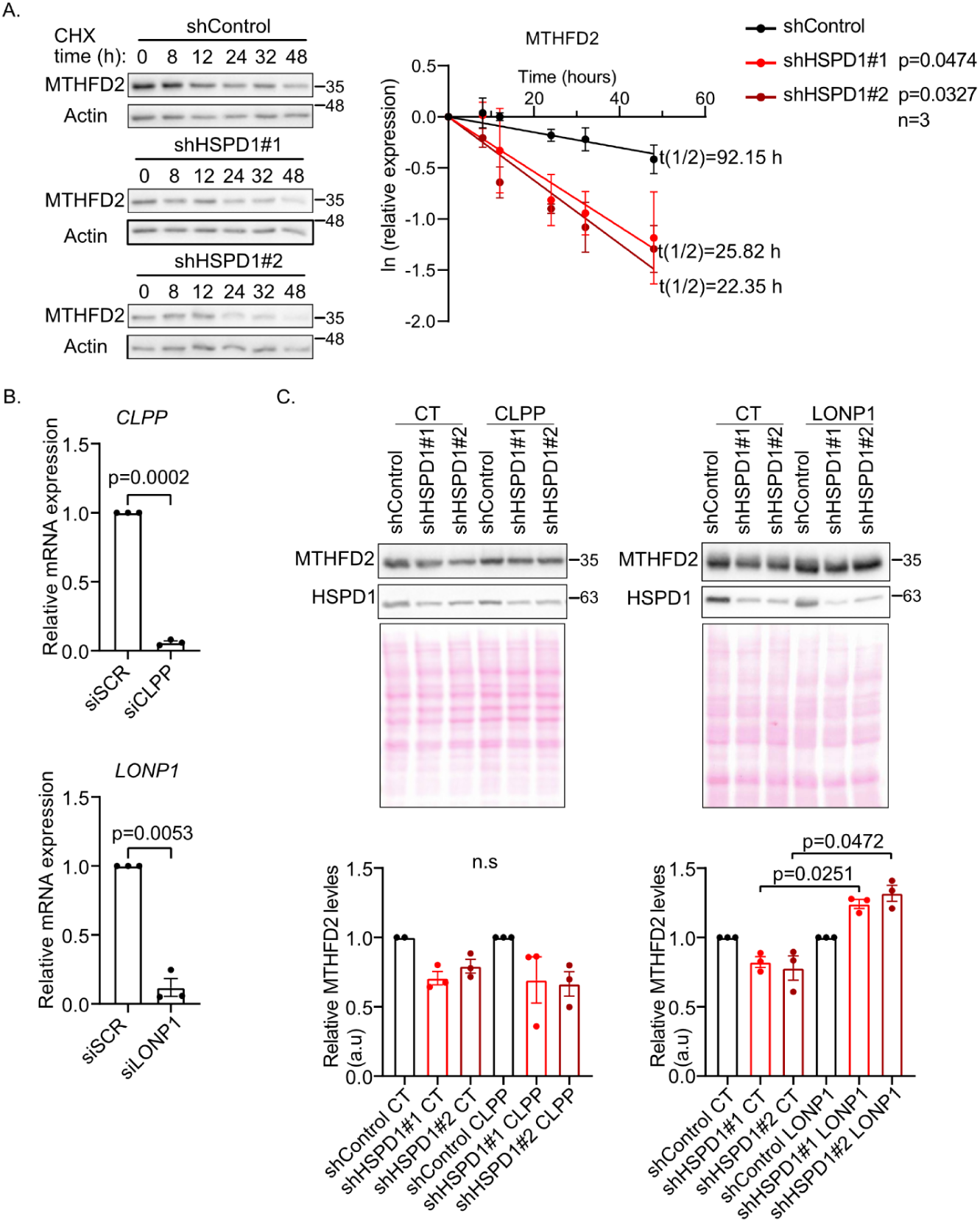
MTHFD2 is degraded by LONP1 in HSPD1 KD cells. A. Cells were treated with CHX for the indicated time points, after which they were lysed and analysed by western blot. Actin was used as a loading control. B. Pooled siRNA were used to KD CLPP and LONP1. KDwas validated using qRT-PCR. C. MTHFD2 levels in siCLPP and siLONP1 in HSPD1 KD and control cells were measured using western blot. Ponceau staining was used as a loading control. Significance was measured in siCLPP and siLONP1 vs. siCONTROL in HSPD1 KD cells.

Mitochondria are equipped with quality control mechanisms. The CLPP and LONP1 proteases have been shown to eliminate misfolded proteins in the mitochondrial matrix^42^. We used siRNA to KD CLPP and LONP1 (Fig. 4b) and used western blot to measure MTHFD2 in HSPD1 KD and control cells (Fig. 4c). LONP1 KD, but not CLPP1 KD, restored MTHFD2 levels in HSPD1 KD cells (Fig. 4d). These results support that MTHFD2 is degraded by LONP1 in HSPD1 KD cells and are in accord with recent data demonstrating that in HSPD1 KD cells, LONP1 degrades aggregation-prone mitochondrial proteins^21^.

### HSPD1 folds MTHFD2 independently of HSPE1

To test whether HSPD1 physically interacts with and folds MTHFD2, we used recombinant MTHFD2 and HSPD1 and a chaperone-assisted folding assay (Fig. 5a). We confirmed that our recombinant MTHFD2 binds CH2methylene-THF using thermal shift assay (Fig. S3a). As a surrogate for proper folding, we measured the enzymatic activity of recombinant MTHFD2 using the substrates methylenemethenyl-THF and NADPH and reading the accumulation of the products, methenylmethylene-THF and NAD, using a plate reader at OD 350 nm (Fig. S3b).

**Fig. 5.**
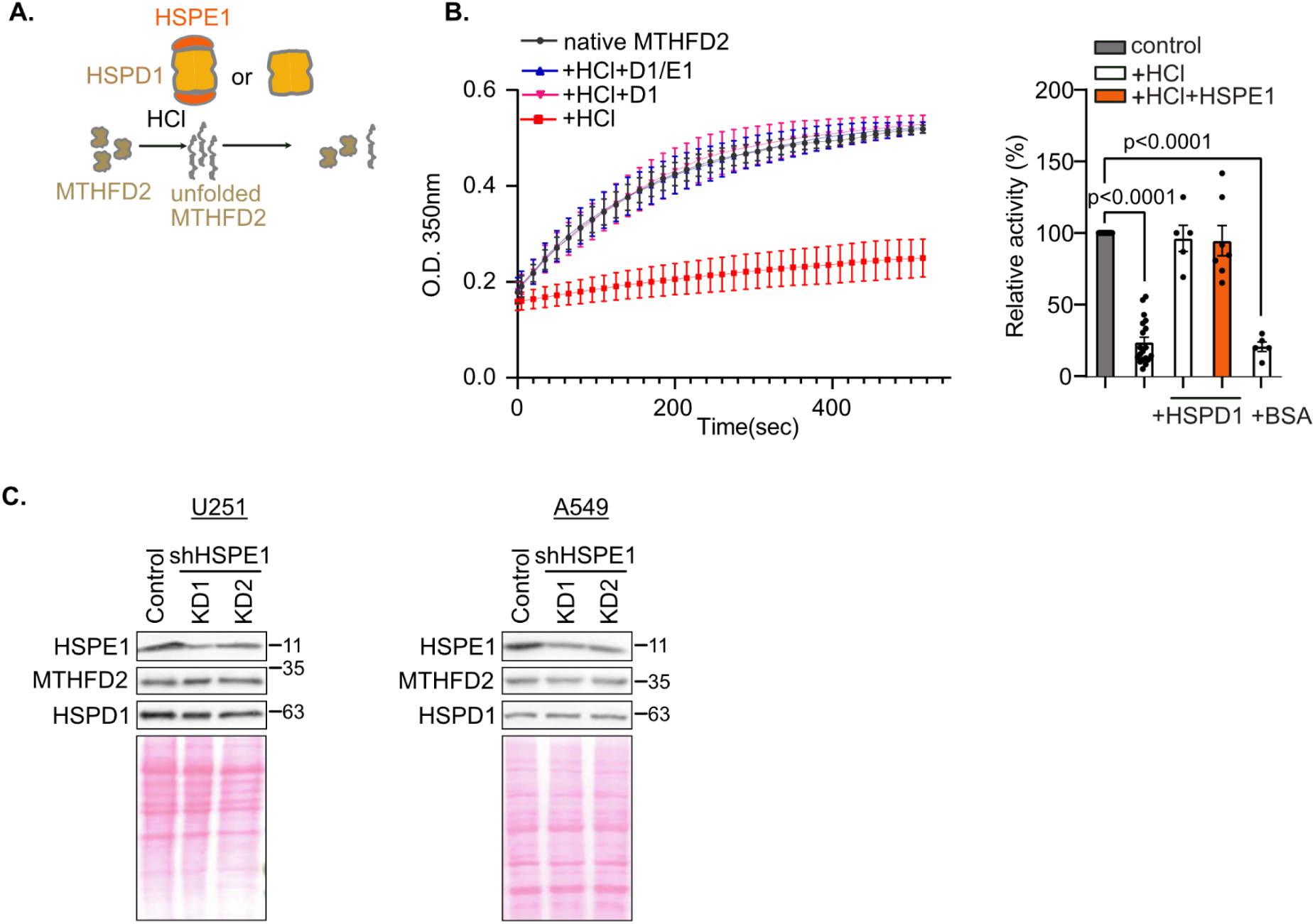
HSPD1 folds MTHFD2 independently of HSPE1. A. Scheme depicting chaperone-assisted folding assay. B. MTHFD2 activity following acid treatment was measured using a plate reader. The slope of the curve was used to determine MTHFD2 activity. BSA was used to control for crowding effect. C. MTHFD2 levels in HSPE1 KD and control cells were measured using western blot. Poncue was used as a loading control.

We treated MTHFD2 with acid to promote unfolding, restored pH to 7.2, and measured MTHFD2 enzymatic activity. We found acid eliminated MTHFD2 enzymatic activity, indicating misfolding. Following acid treatment and neutralization, we added HSPD1 and HSPE1 to the misfolded MTHFD2 and immediately measured MTHFD2 activity. We found that HSPD1 restored MTHFD2 to its fully active form (Fig. 5b). Surprisingly, HSPD1 could do so independently of its co-chaperone, HSPE1 (Fig. 5b). Interestingly, it was previously reported that the bacterial homologue of HSPD1, GroEL, folds misfolded bovine rhodanese independently of GroES, the bacterial homologue of HSPE1^43^.

Structural studies of the HSPD1-HSPE1 complex and their closely related bacteria homologs, GroEL-GroES, support that HSPE1 and GroES play key roles in HSPD1 and GroEL substrate folding activity^44^. Having found that HSPD1 folds MTHFD2 independently of HSPE1 *in vitro* (using recombinant proteins), we predicted that HSPD1 would be sufficient to support MTHFD2 expression in cells. We used shRNA to KD HSPE1 in A549 cells and measured MTHFD2 levels in HSPE1 KD and control cells (Fig. 5c). We found that HSPE1 depletion did not affect MTHFD2 or HSPD1 expression levels.

To identify the role of HSPE1 in mitochondrial respiration, we used Seahorse XP and found that HSPE1 KD leads to reduced basal respiration but not maximal respiration (Sup 3c). In addition, HSPE1 did not affect mitochondrial mass in A549 cells (Sup 3d). Together, these results suggest that HSPE1 KD does not lead to total mitochondrial collapse and that HSPE1 does not play an important role in MTHFD2 folding.

### Divergent transcriptional response to HSPD1 and HSPE1 KD

Since we found HSPD1 folds MTHFD2 independently of HSPE1 (Fig. 5) and since ecological network analysis using gene expression obtained from different tumor types showed that HSPD1 and its co-chaperone HSPE1 are co-expressed with a different set of mitochondrial proteins^14^, we hypothesized that HSPD1 and HSPE1 cater to the folding of different proteins and play divergent biological functions. We reasoned that if the major biological functions of HSPD1 and HSPE1 stem from their interaction as a fully formed chaperonin and are both required to fold the same set of proteins, similar transcriptional changes will be observed following KD of either. However, if the major biological functions of HSPD1 and HSPE1 are independent, we predict that the transcriptional response to KD of either will be very different.

We compared the cellular response to HSPD1 and HSPE1 KD using RNAseq, and in line with the latter, we found vast differences in the transcriptome of HSPD1 KD and HSPE1 KD A549 cells (Fig. 6a; Table S2). More than 700 transcripts were up- or down-regulated following HSPD1 or HSPE1 KD, of which <10% were shared between the chaperone and its co-chaperone (Fig. 6b, Table S2). These differences were also reflected in the biological categories enriched in the list of up- or down-regulated transcripts in HSPD1 and HSPE1 KD cells (Fig. 6c). For example, categories related to oxidative stress, such as glutathione metabolic processes, were enriched in the list of down-regulated transcripts in HSPD1 KD but not HSPE1 KD cells (Fig. 6c). Conversely, categories related to RNA processing were enriched in the list of down-regulated transcripts in HSPE1 KD but not HSPD1 KD cells (Fig. 6c).

**Fig. 6.**
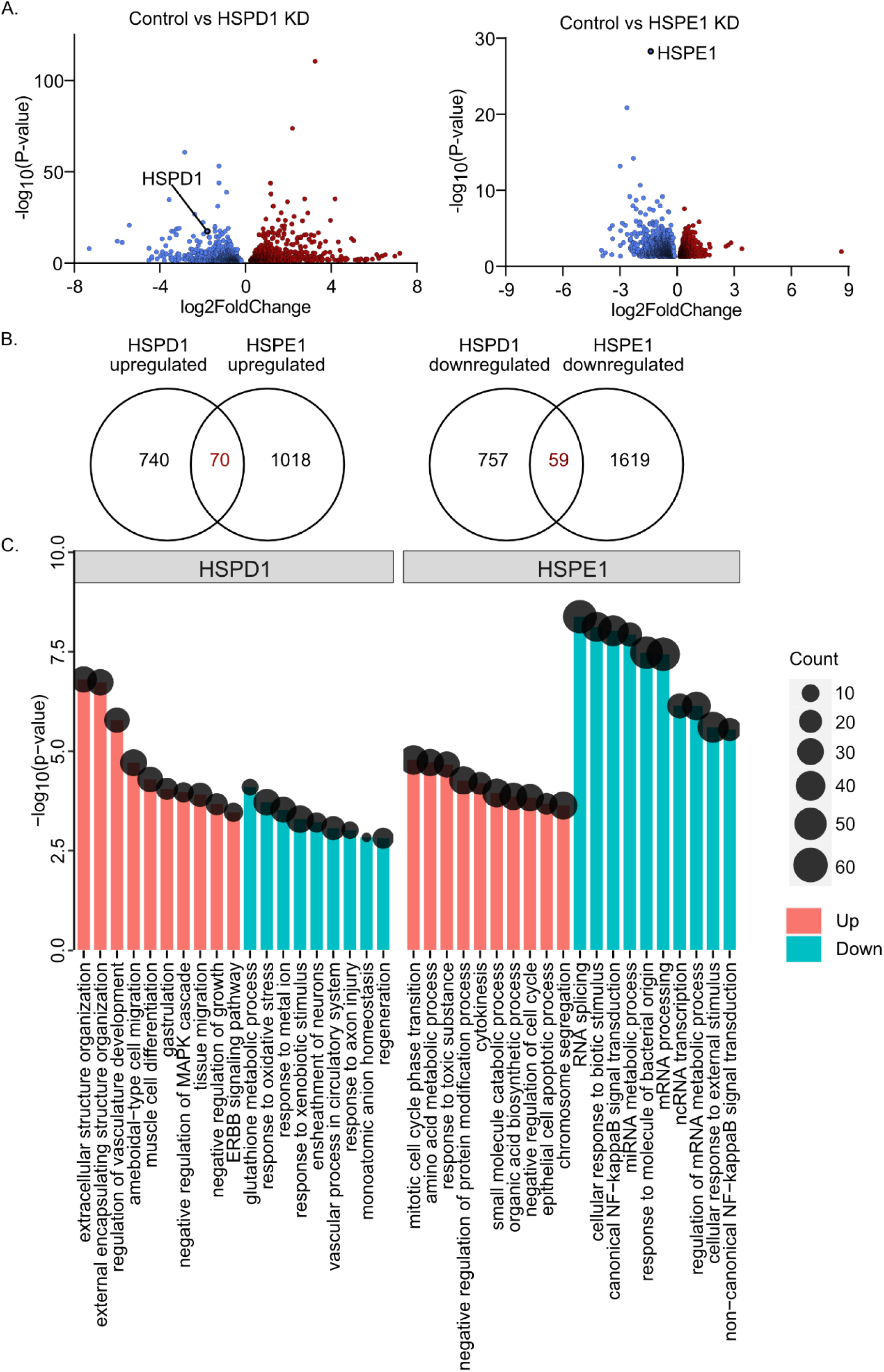
The impact of HSPD1 vs. HSPE1 KD on gene expression. A. Gene expression in HSPD1 or HSPE1 KD vs. control U251 cells was measured using RNAseq (3 biological replicates per condition; pooling the two KD cell lines together) and is shown using a volcano plot. B. Ven diagram of up- or down-regulated transcripts in HSPD1 KD and HSPE1 KD cells. C. Pathways enriched in transcripts upregulated or downregulated by HSPD1 and HSPE1 KD.

We validated these findings in U251 cells (Fig. S4a, Table S2) and found that in these cells as well, there were profound transcriptional changes following HSPD1 or HSPE1 KD. More than 700 transcripts were affected by the KD, with an overlap of 17 transcripts whose expression changed by either HSPD1 or HSPE1 KD (Fig. s4b). In line with the results obtained using A549 cells, the biological categories enriched in the list of up- or down-regulated transcripts in HSPD1 and HSPE1 KD cells were very different. For example, transcripts related to actin filament organization were upregulated in HSPD1 KD cells, while cytoplasmic translation was upregulated in HSPE1 KD cells (Fig. S4). Taking these results together, we conclude that the transcriptional response to downregulation of HSPD1 or HSPE1 is cell line-specific. In addition, these data show that the transcriptional response to KD of either HSPD1 or HSPE1 is very different, in line with the hypothesis that they cater to the folding of distinct groups of proteins.

### Divergent metabolic response to HSPD1 and HSPE1 KD

Having found divergent transcriptional changes following KD of HSPD1 vs. HSPE1, we asked if such a divergent response is reflected in the cellular metabolic profile. To this end, we used targeted metabolomics analysis to identify metabolic changes in response to HSPD1 or HSPE1 KD in A549 cells growing in human plasma-like media. We designated a positive hit to metabolites that were significantly decreased or increased in the two KD cell lines vs. control cells (p<0.05 following multiple test correction). Of 216 metabolites analyzed, two were down-regulated and four up-regulated in HSPD1 KD cells (Fig. 7a; Table S3-5). In HSPE1 KD vs. control cells, 10 metabolites were upregulated (Fig. 7c). Two metabolites were upregulated in both HSPD1 and HSPE1 KD cells (Fig. 7b).

**Fig. 7.**
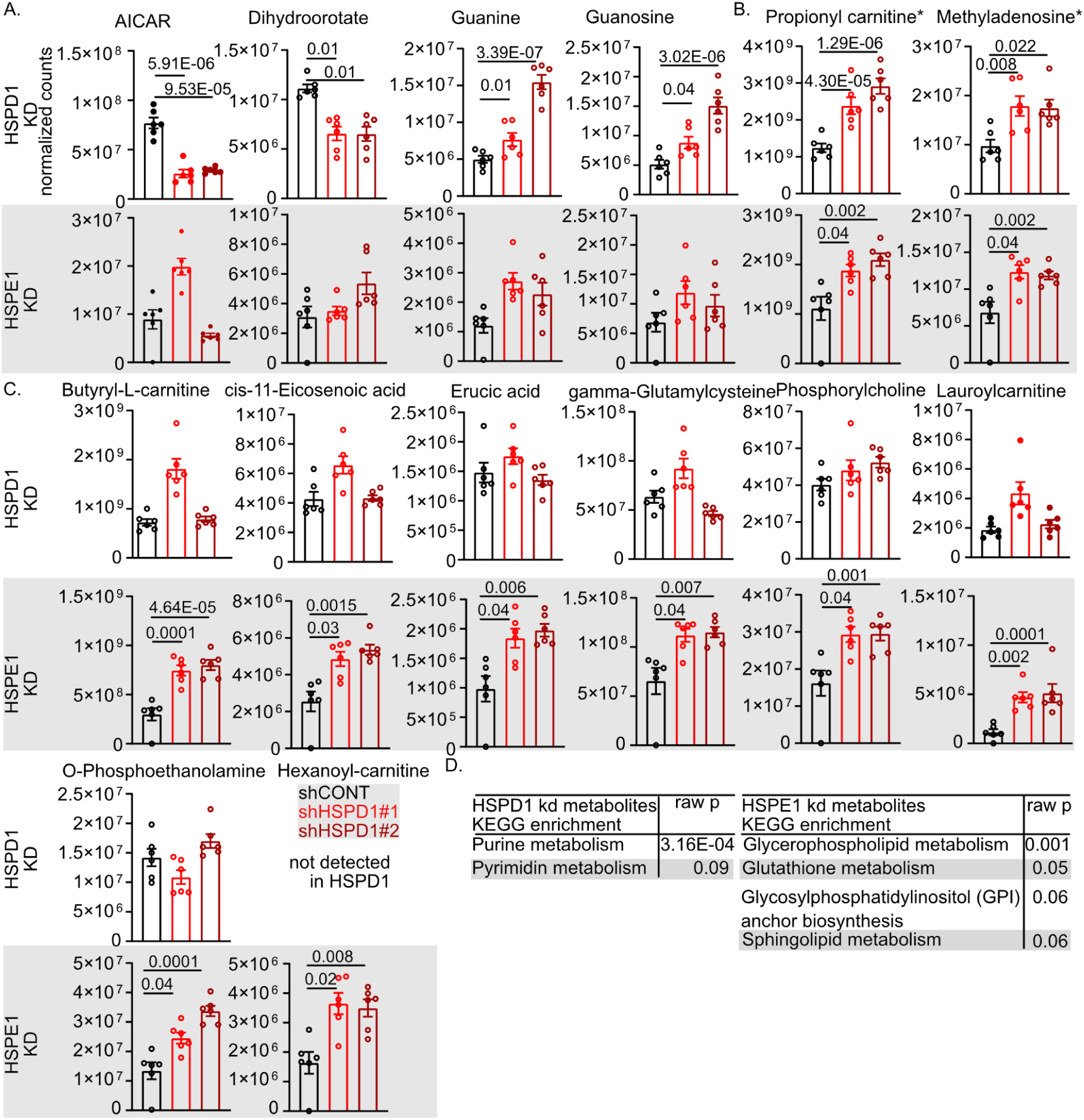
Metabolites in HSPD1 KD and control cells. A. Metabolites significantly affected by HSPD1 KD (same direction in two KD cell lines) but not changed by HSPE1 KD. B. Metabolites significantly affected in both HSPD1 and HSPE1 cells. C. Metabolites affected by HSPE1 KD but not by HSPD1 KD. D. MetaboAnalyst identified enriched metabolic pathways in HSPD1 and HSPE1 KD cells.

We used Metaboanalyst 6.0 (metaboanalyst.ca) to identify pathways enriched in our list of metabolites significantly affected by HSPD1 and HSPE1 depletion (Fig. 7D). Purine Metabolism was the most significantly enriched pathway in HSPD1 KD cells, which is directly linked to the 1C pathway^36^. The reduced AICAR found in HSPD1 KD cells suggests that HSPD1 not only regulates MTHFD2, as MTHFD2 depletion alone leads to AICAR accumulation^45^, and is compatible with reduced MTHFD2 along with other components of the 1C pathway. For example, decreased ACAR levels were found in cells depleted of MTHFD2 and PAICS^46^. In contrast, Glycerophospholipid Metabolism was found enriched in HSPE1 KD cells (Fig. 7c). Interestingly, the accumulation of acylcarnitines (Butyryl-L-carnitine, Hexanoyl carnitine, Lauroylcarnitine, Propionyl carnitine) in HSPE1 KD cells suggests compromised mitochondrial β-oxidation^47^, which is in line with previous metabolic analysis of the consequence of HSPE1 depletion in neurons^48^. These results show that cells acquire distinct metabolic states following depletion of HSPD1 vs. HSPE1, further supporting that the two have divergent biological functions.

### The mitochondrial stress response is activated in different tissues following HSP60 or HSP10 knockout in C. elegans

Our proteomic, transcriptomic, and metabolic data support that HSPD1 and HSPE1 have divergent functions. To test this hypothesis in a whole organism, we used C. elegans. We employed CRISPR KO of either HSP60 or HSP10. The two KO strains caused larval arrest at the L2-L3 stage and were thus maintained as heterozygotes.

To test if HSP60 and HSP10 KO have divergent functions, we focused on the activation of the Mitochondrial Unfolded Protein Response (UPRmt) as a readout of insufficient mitochondrial proteostasis. To this end, we crossed worms engineered to harbor GFP under the *hsp-6* (*C. elegans* HSPA9 ortholog) promoter, which is activated by ATFS-1, a classic readout of the Mitochondrial Unfolded Protein Response^49,50^, to our KO heterozygote strains. We chose this model because we assume that if KO of *hsp-60* or *hsp-10* results in a differential mitochondrial impact, we will observe distinct GFP expression patterns. In other words, if in general, HSP-60 and HSP-10 were to work together to fold the same subset of proteins, we would expect that similar tissues will experience UPRmt, and a similar GFP expression pattern will be observed in the case of either *hsp-60* or *hsp-10* KO. We found that *hsp-10* KO mutants exhibited GFP expression in muscles, while *hsp-60* mutants expressed GFP in the gut (Fig. 8A). In accord, we found decreased motility rates in *hsp-10* but not in *hsp-60* KO worms (Fig. 8B). These data support that HSP-60 and HSP-10 have divergent functions in the whole organism and that such divergent functions are found in nematodes as well as in cancer cell lines.

**Fig. 8.**
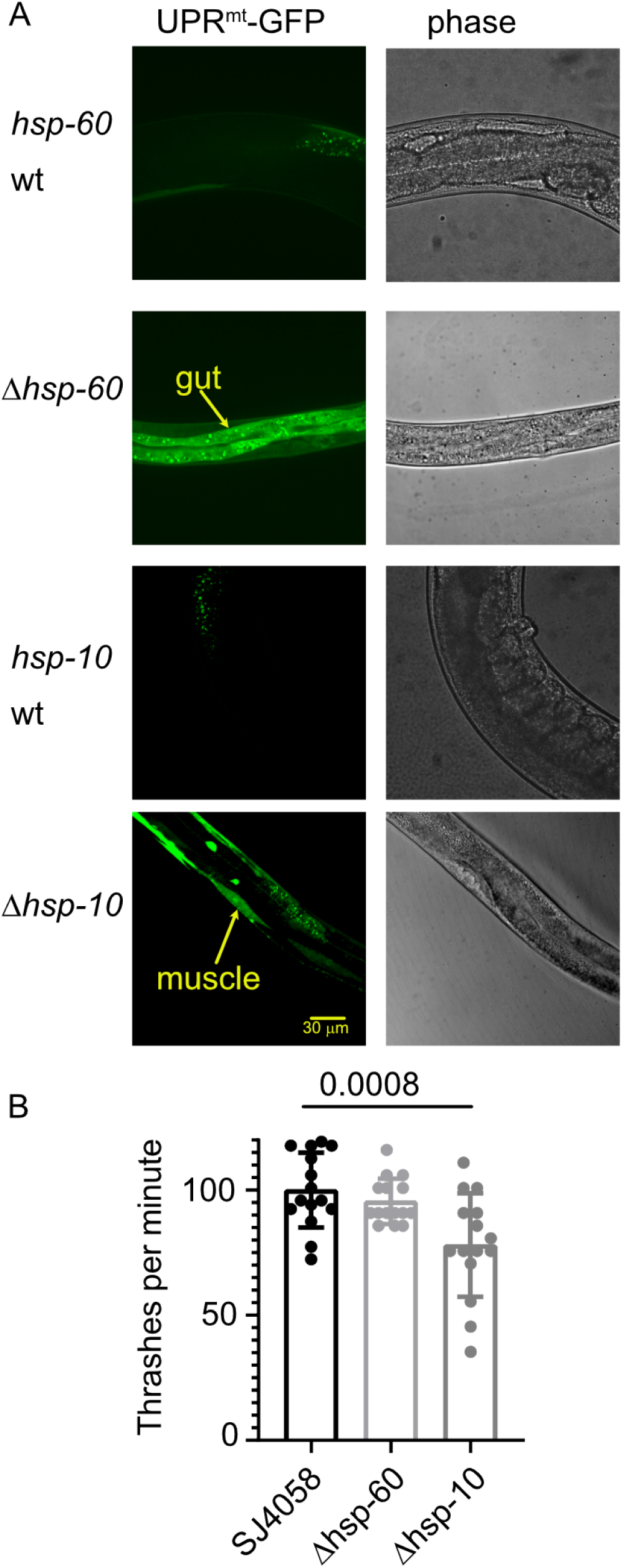
Divergent phenotypes of Hsp60 vs. Hsp10 KO in C. elegans. A. GFP expression indicates UPRmt activation in the gut of hsp-60 KO and in the muscles of hsp-10 KO worms. B. Thrashing was recorded in the indicated strains.

## Discussion

Using SILAC-based proteomics and HSPD1 KD cells, we found that MTHFD2 protein expression depends on HSPD1. We confirmed that MTHFD2 is an endogenous client of HSPD1, that HSPD1 folds MTHFD2 independently of HSPE1 using recombinant proteins, and that tumor and C. elegans cells respond very differently to HSPD1 vs. HSPE1 depletion. We will discuss the implications of these findings for understanding the role of HSPD1 in supporting cellular metabolism, the mitochondrial chaperone-client network, and the HSPE1-independent role of HSPD1 as a molecular chaperone.

Previously, HSP60 and HSP10 temperature-sensitive yeast mutants were used to identify HSP60 and HSP10 client proteins in isolated mitochondria^51^. Nine proteins were identified and validated as potential HSP60 or HSP10 clients, none of which were identified in our SILAC experiment. HSPD1 mutants associated with human pathologies have been used to identify HSPD1 clients and to infer possible mechanisms linking HSPD1 mutations to the misfolding of HSPD1 substrates and diseas^52,53^. More than 400 putative clients were identified, 60 of which were identified in our SILAC experiment using HSPD1 KD cells, including the mitochondrial 1C enzyme MTHFD2^5^. In support, analysis of multiple knockout mouse models experiencing severe oxidative phosphorylation (OXPHOS) dysfunction revealed upregulation of HSPD1 and MTHFD2 protein and transcript levels as a consistent and early response to mitochondrial stres^54^. Recently, HSPD1 and HSPE1 were shown to play important roles in mitochondrial translation by promoting mitochondrial ribosomal proteins ^21^ and a mitochondrial import proteins ^24^. Nevertheless, whether the previously identified proteins are truly endogenous HSPD1 or HSPE1 clients, or whether HSPD1 can directly fold these proteins, with or without HSPE1, is not known. Here, we used a gene KD approach, degradation assays, and an *in vitro* chaperon-assisted folding assay to show that MTHFD2 is an endogenous HSPD1 client.

The mitochondrial 1C pathway in general, and MTHFD2 in particular, is critical for NADPH and nucleotide generation ^40,55^. In addition, MTHFD2 is upregulated by oncogenic signaling, including mTOR activation and MYCN amplification in neuroblastoma, and is essential for metabolic reprogramming induced by these pro-oncogenic signaling proteins^35,37^. It is therefore not surprising that MTHFD2 is a drug target in cancer ^56,57^. We found that HSPD1 is co-expressed with MTHFD2, essential for maintaining MTHFD2 levels, and that HSPD1 KD leads to reduced SAM levels and increased ROS, suggesting that HSPD1 promotes the 1C pathway by folding MTHFD2 and that targeting HSPD1 will be impactful in cases where reduced 1C output is clinically desirable.

The mitochondrial chaperone-client co-expression network in cancer reveals that HSPD1 and HSPE1 cluster in distinct groups, despite co-expression and sharing a common promoter^14^. One possible explanation for this finding is that HSPE1 is essential for the folding of a subset of HSPD1 clients and has evolved to be co-expressed with proteins that depend on it, but not with those that do not. A reasonable prediction from this model is that cells will respond differently to HSPD1 vs. HSPE1 depletion. Using RNAseq and metabolomics, we found little overlap in the transcriptional and metabolic responses of cells to KD of either HSPD1 or HSPE1, suggesting that they have divergent biological functions. Moreover, we found that UPRmt activation occurred in distinct tissues in response to KO of either HSP60 or HSP10 in *C. elegans*. If HSP10 were to function only as a co-chaperone helping HSP60 fold some of its clients, we would expect to find that activation of UPRmt in the HSP10 KO tissue would be nested in the UPRmt activated HSP60 KO tissue. In contrast, we found that HSP10 exhibited a unique pattern of UPRmt, where specific activation of UPRmt in muscle not observed in the HSP60 KO. These data support that HSP60 and HSP10 evolved significantly divergent functions.

It was previously shown that GroEL can fold an exogenous target independently of GroES and ATP, acting as a catalytic unfoldase^43^. It was also found that HSPD1 interacts with ClpP through its epical domain to promote mitochondrial functions and tumorogenicity in cancer cells^58^. Using MTHFD2 and recombinant HSPD1, we found that HSPD1 can fold an endogenous client independently of HSPE1. In accord, HSPD1 and HSPE1 fractionated differently in thermal proteome profiling following mitochondrial stress activation^59^, indicating that they interact with distinct protein complexes in these conditions.

In summary, we found that HSPD1 is an MTHFD2 chaperone and that HSPD1 and HSPE1 have divergent functions, raising the possibility that HSPE1 has important cellular functions beyond acting as an HSPD1 co-chaperone.

## Methods

In supplementary material

## Supporting information

sup data, methods, full scan blots

table s1-SILAC

table s2-rnaSEQ

table s3-metabolomics HSPD1

table s4-metabolomics HSPE1

table s5-metabolomics

## Acknowledgments

Research reported in this publication was supported by the Israel Science Foundation (grant number 228/25 BR and 420/23 ABZ) and by Worldwide Cancer Research (grant reference number 24-0072) for BR.

