## Supplementary material for "The mitochondrial chaperone HSPD1 folds MTHFD2 independently of its co-chaperone HSPE1": sup data, methods, full scan blots

* equal contribution

### Corosponding

AFFILIATIONS

1 Department of Life Sciences, Ben-Gurion University of the Negev, Beer-Sheva, Israel

2 Ilse Katz Institute (IKI) for Nanoscale Science and Technology, Ben-Gurion University of the Negev, Beer Sheva, Israel

3 Laura and Isaac Perlmutter Metabolomics Center, B. Rappaport Faculty of Medicine, Technion-Israel Institute of Technology, Haifa, Israel.

4 The Shraga Segal Department of Microbiology, Immunology and Genetics, Faculty of Health Science, Ben-Gurion University of the Negev, Beer-Sheva, 84105, Israel.

5 Faculty of Health Sciences, Ben-Gurion University of the Negev, Beer-Sheva, 84105, Israel.

Fig S1

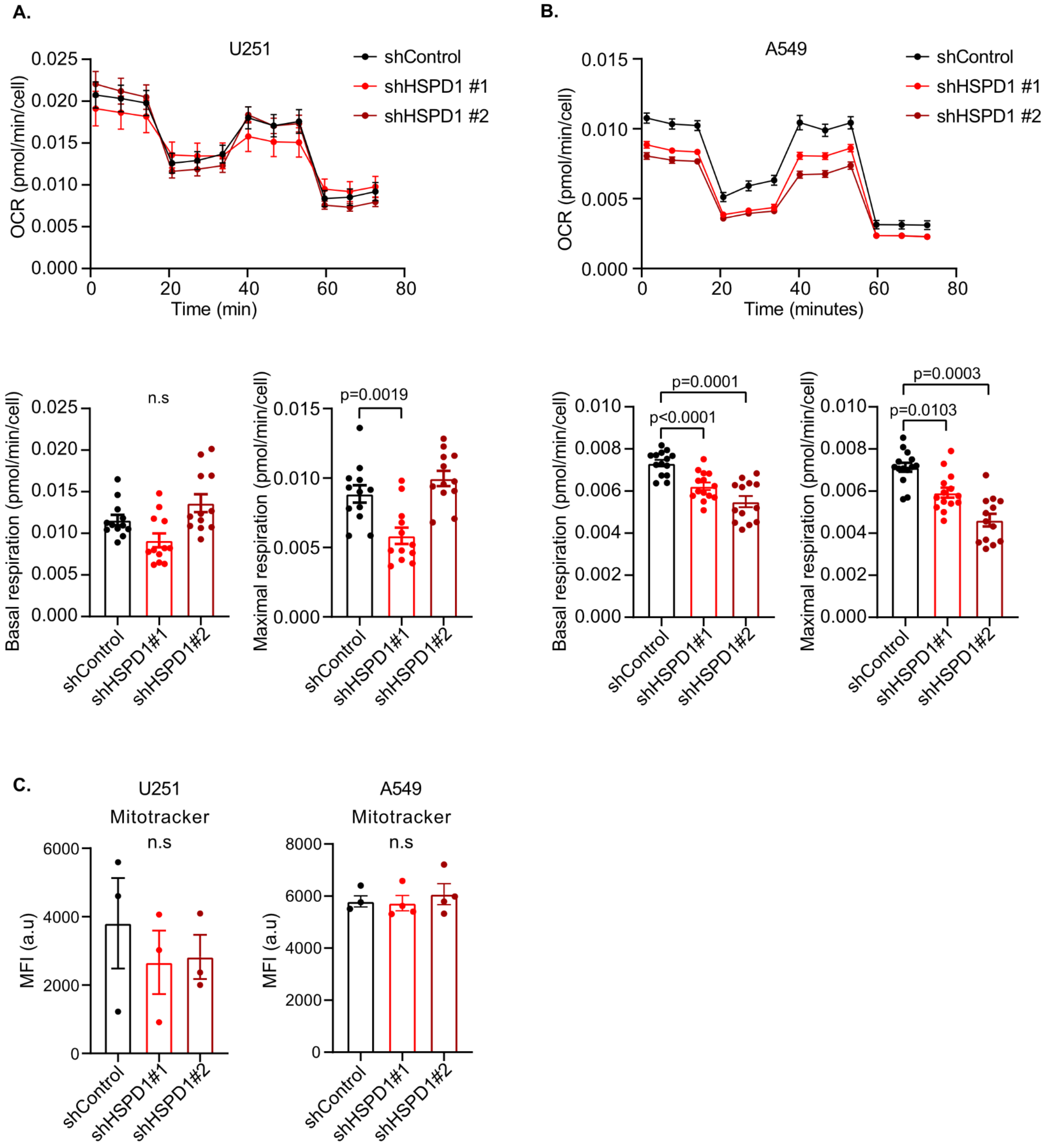


**SUP Fig. 1.**

A, B. Seahorse XF analysis of mitochondrial function.

C. Mitochondrial mass in HSPD1 KD and control cells was measured using Mitotracker Green and FACS.

Sup 2

A.

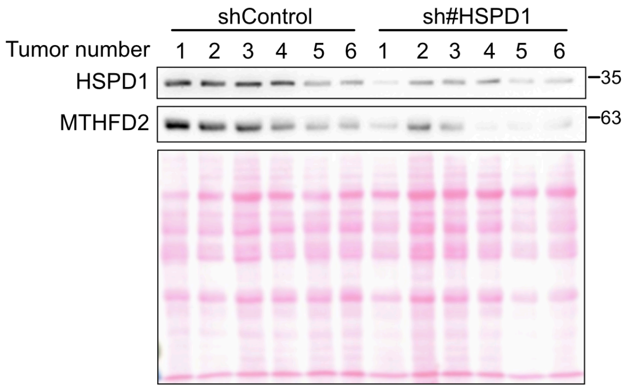


**SUP Fig. 2.**

A. Lysates from HSPD1 kd and control tumors were analysed by western blot. Poncue was used as a loading control.

Sup fig. 3

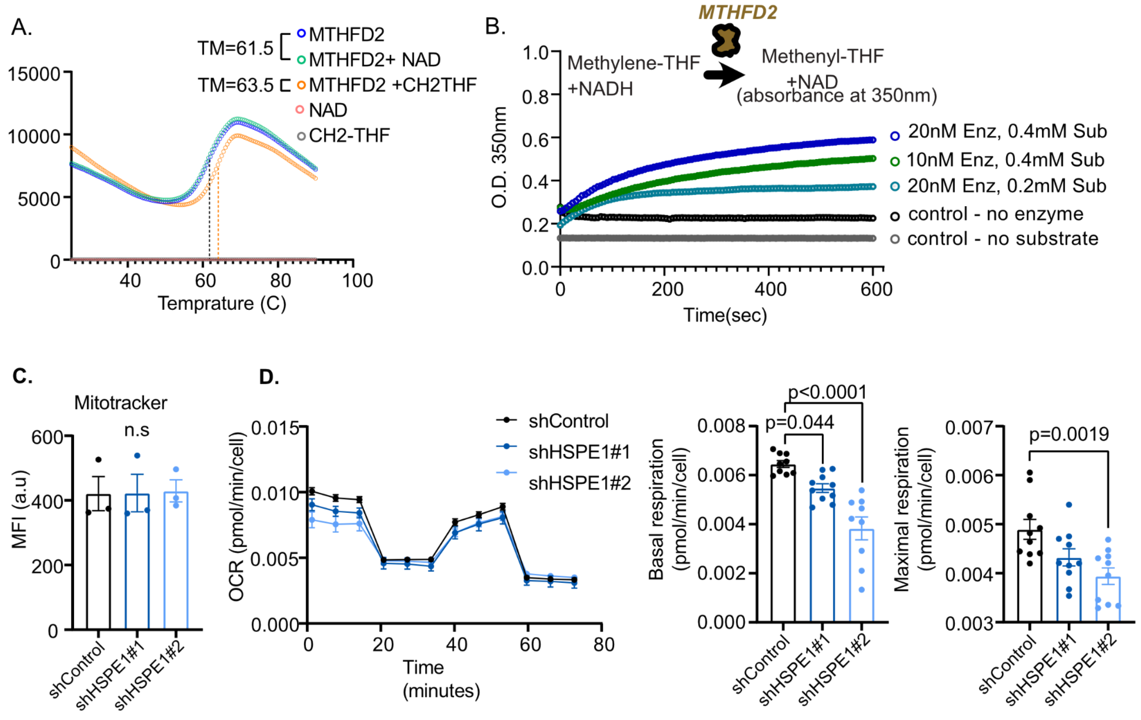


**SUP Fig. 3.**

A. Thermal shift assay using recombinant MTHFD2 and CH2THF.

B. MTHFD2 activity assay using a plate reader.

C. Mitochondrial mass HSPE1 KD and control cells were measured using Mitotracker Green and FACS.

D. Seahorse PX was used to measure mitochondrial activity in HSPE1 kd and control cells.

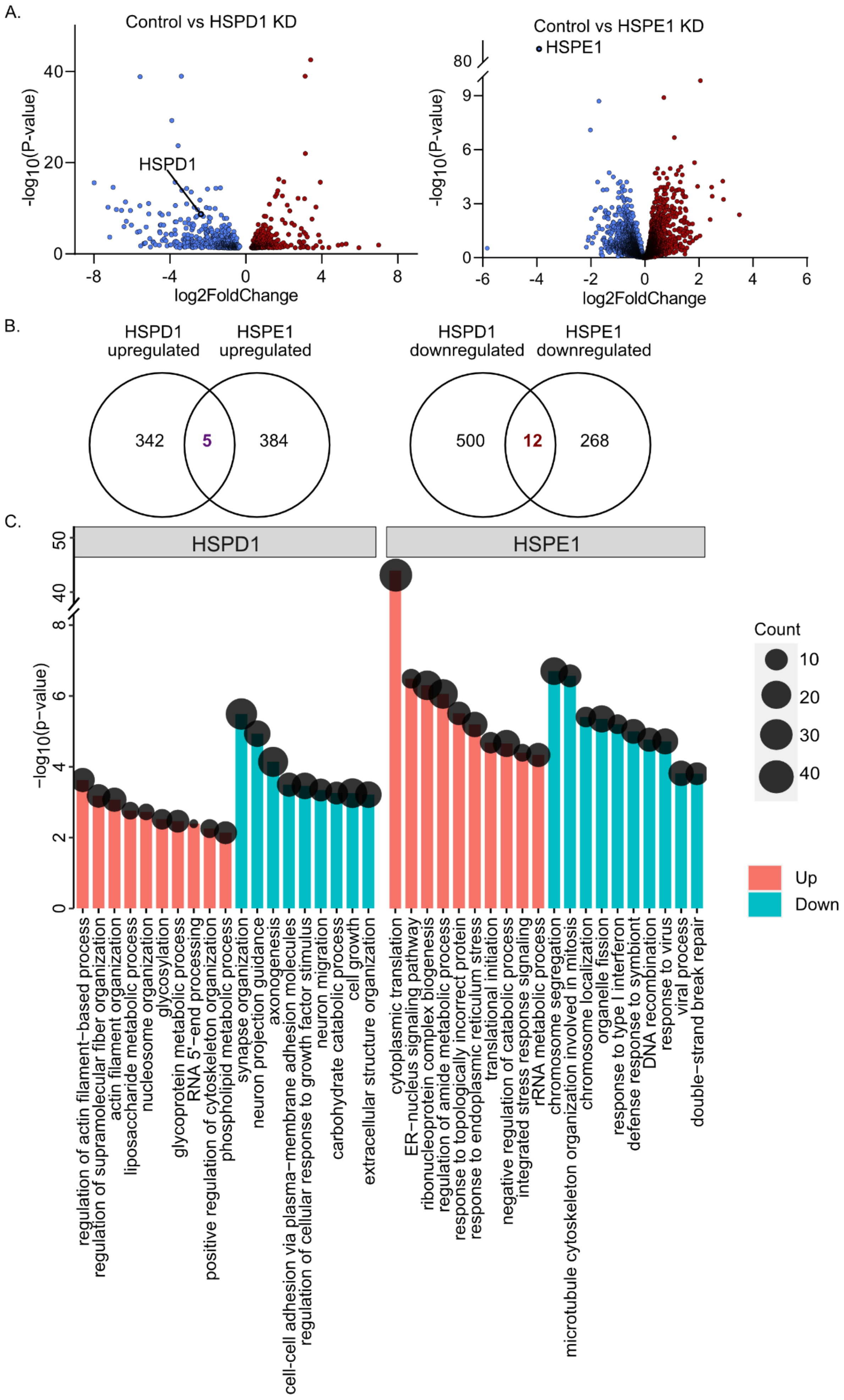
A.
Control vs HSPD1 KD
-log10(P-value)
40
20
0
HSPD1
-8
-4
0
4
8
log2FoldChange
Control vs HSPE1 KD
HSPE1
80
9
6
3
0
-6
-4
-2
0
2
4
6
log2FoldChange
B.
HSPD1 upregulated
HSPE1 upregulated
342
5
384
HSPD1 downregulated
HSPE1 downregulated
500
12
268
C.
HSPD1
HSPE1
−log10(p−value)
50
40
8
6
4
2
0
Count
10
20
30
40
Up
Down
regulation of actin filament−based process
regulation of supramolecular fiber organization
actin filament organization
liposaccharide metabolic process
nucleosome organization
glycosylation
glycoprotein metabolic process
RNA 5'−end processing
positive regulation of cytoskeleton organization
phospholipid metabolic process
synapse organization
neuron projection guidance
axonogenesis
cell-cell adhesion via plasma−membrane adhesion molecules
regulation of cellular response to growth factor stimulus
neuron migration
carbohydrate catabolic process
cell growth
extracellular structure organization
cytoplasmic translation
ER−nucleus signaling pathway
ribonucleoprotein complex biogenesis
regulation of amide metabolic process
response to topologically incorrect protein
response to endoplasmic reticulum stress
translational initiation
negative regulation of catabolic process
integrated stress response signaling
rRNA metabolic process
chromosome segregation
microtubule cytoskeleton organization involved in mitosis
chromosome localization
organelle fission
response to type I interferon
defense response to symbiont
DNA recombination
response to virus
viral process
double−strand break repair

**SUP Fig. 4.**

A. Gene expression in HSPD1 or HSPE1 KDbvs. control cells was measured using RNAseq (4 biological replicates per condition; pooling the two kd cell lines together) and are shown using a volcano plot.

B. Ven diagram of up- or down-regulated transcripts in HSPD1 KD and HSPE1 KD cells.

C. Pathways enriched in transcripts upregulated or down-regulated by HSPD1 and HSPE1 KD.

### Methods

#### Cell culture

Human breast cancer cell line MDB_MB_231, human glioblastoma cell line U251, lung cancer cell line A549, and ovarian cancer OAW42 were obtained from ATCC. All cell lines were cultured in Dulbecco's Modified Eagle Medium (DMEM) Medium was supplemented with 10% fetal bovine serum (FBS), sodium pyruvate solution (1mM), and antibiotic–antimycotic (complete DMEM). Cells were kept in an incubator with 5% $CO_2$ at 37°C.

#### Generation of stable cell lines for gene knockdown

Gene knockout of HSPD1 was performed using the SYNTHEGO Gene Knockout Kit v2. A549 and U251 cells were seeded into 6-well plates at a density of 200,000 cells per well and allowed to grow overnight. The next day, cells were transiently transfected with HSPD1-specific CRISPR guide RNAs pool (gRNAs) and CAS9 to a final concentration of 3uM, using Lipofectamine Cas9 Plus reagent and following the recommended protocol. HSPD1 protein levels were measured using western blot three days post-transfection, and three “pool-CRISPR-knockdown” populations were analyzed. WT cells were transfected similarly, without CAS9. Next, a single clone isolation protocol was performed to select clones with HSPD1 downregulation. Pool-CRISPR cells were seeded in 96 well plates at a concentration of 0.5 cells well. Filtered DMEM from proliferating U251 WT cells was used in addition to fresh DMEM until small colonies were observed by light microscopy. Once clones filled a well, they were transferred to a larger well plate. Immunoblotting was used to validate the HSPD1 knockdown.

#### Lentiviral silencing

Two different CRISPR clones were chosen for HSPD1 knockdown using two independent (Short hairpin RNA) shRNA (SIGMA ALDRICH, MISSION shRNA, TRCN0000029446 and TRCN0000343952 for sh_1 and sh_2, respectively). For lenti virus generation, HEK293T Cells were grown in complete DMEM and transfected using CalFectin and a plasmid ratio of 1:2:3 (PAX2; pMD2.G; transfer vector). The media was changed after 24 hours, and the viruses were collected after 48 hours. The collected viruses were stored at -80°C. Prior to virus infection, cells were grown to 60% confluence on 6-well plates and then infected with the virus at a ratio of 1:10 (shCONT, sh#1, sh#2) and left for 24 hours in a 37°C incubator. Following infection, the containing medium was removed, replaced with a fresh medium, and incubated at 37°C. Puromycin (1:1000) was used for selecting infected cells. Gene knockdown was validated using qRT-PCR and Western blot.

#### siRNA transfection

Knockdown of CLPP or LONP1 was achieved using siRNA pools specific to each target, while a scrambled siRNA (siSCR) was used as a control (Horizon). Transfection was carried out using the LipoJet Transfection Kit (SignaGen Laboratories) according to the manufacturer's protocol. Briefly, cells were seeded in a 6-well plate one day before transfection and allowed to grow to approximately 60% confluence. The desired concentration of each siRNA was prepared (final concentrations- siSCR: 50 nM or 80 nM, siRNA for CLPP: 50 nM, siRNA for LONP1: 80 nM) and added to the cells along with LipoJet. After 48 hours, the cells were harvested for Western blot analysis or qPCR.

#### Quantitative Reverse Transcription Polymerase Chain Reaction (qRT-PCR).

Total RNA was extracted from cells using an RNA purification kit (Invitrogen), following the manufacturer's instructions. RNA concentration and quality were determined using a NanoDrop spectrophotometer. cDNA synthesis was performed using the cDNA synthesis kit (BioLabs), according to the manufacturer's protocol, with approximately 800 ng of RNA as the input. For qPCR, reactions were carried out in a final volume of 12 μL, containing 6 μL of 2× SYBR Green Master Mix (PCR Biosystems), 1 μL of cDNA, 0.5 μM of forward and reverse primers (purchased from IDT), and 4 μL of nuclease-free water. PCR reactions were performed on iCycler Real-Time PCR System (Bio-Rad Laboratories) with the following cycling conditions: initial denaturation at 95°C for 10 minutes, followed by 46 cycles of 95°C for 10 seconds, 58°C for 30 seconds, and 72°C for 3 seconds. Finally, an elongation step was performed at 40°C for 10 minutes to ensure complete extension of any remaining single-stranded DNA. qRT-PCR data were acquired using the iCycler software, with threshold cycle (Ct) values determined from 4 replicate reactions. Relative gene expression was calculated using the 2^(-ΔΔCt) method, normalized to the expression of L32 as a housekeeping gene.

#### Western blot

Cells were cultured in a 10 cm dish to 90%-95% confluency and were lysed with RIPA lysis buffer supplemented with protease inhibitor (1:10000) and phosphatase inhibitor (1:10000). The lysates were sonicated for homogenization, and Pierce™ BCA Protein Assay Kit was used for protein quantification. A minimum of 10 μg of the protein lysate was subjected to sodium dodecyl sulfate-polyacrylamide gel electrophoresis (SDS- PAGE) and transferred onto nitrocellulose (BioTrace™, #66485) membranes. Membranes were stained with ponceau S and filmed before blocking with 5% skim milk powder in Tris buffer saline (TBS) for 1 hour and incubated with primary antibody diluted in blocking buffer overnight at 4°C. Next, the membranes were washed with TBS and incubated with HRP-linked secondary antibodies. Signals were determined using a chemiluminescence detection kit (Advansta, #K-12043-D20). ImageJ software was used to quantify band intensities.

#### Proximity ligation assay and mitochondria staining

Cells were seeded on coverslips one day before the staining. WT A549 cells were stained explicitly for mitochondria labeling utilizing MitoTracker. Briefly, the cells were incubated with a final concentration of 200nM MitoTracker™ Red CMXRos reagent in DMEM for 20 minutes within the incubator. The MitoTracker reagent was removed, and the cells were washed twice with PBS. Proximity Ligation Assay (PLA) was carried out for WT and HSPD1 KD cells, with or without MitoTracker staining, and tumor tissues obtained from mice xenograft model. Before the PLA, the cells/tissued were fixed with 4% paraformaldehyde (PFA) in PBS washed with either PBS (for cells) or PBST (for tissues), and cells were permeabilized with 0.5% of X100 triton in PBS. PLA was conducted using Duolink in situ reagents (Sigma-Aldrich) according to the manufacturer's protocol. In brief, cell or tissue samples were blocked with blocking solution for 30 minutes at room temperature and co-stained with primary antibodies (targeting HSPD1 and MTHFD2) in antibody diluent overnight at 4°c. After this incubation, samples were washed twice with buffer A, and PLA probes (secondary antibodies conjugated oligonucleotides) in antibody diluent were incubated with the cells for 60 minutes in a humidified chamber at 37°c followed by two washes with buffer A. Next, ligase in ligation buffer was incubated with samples for 30 minutes in a humidified chamber at 37°c. This step allows the ligation of the oligonucleotides if their proximity is ≤40nm. After two washes with buffer A, a signal amplification step was carried out by incubating the samples with polymerase diluted with amplification buffer in a humidified chamber at 37°c for 100 minutes. Finally, the samples were washed twice with buffer B and once with 0.1% buffer B. The coverslips were then mounted onto slides using a DAPI-containing mounting medium (SouthernBiotech) and sealed with nail polish to ensure fixation. Images were acquired using laser scanning microscopy (LSM) and spinning disk confocal microscopes.

#### Cycloheximide Chase Analysis

Cells were seeded a day before the experiment, followed by the incubation with a final concentration of 10μM cycloheximide (CHX) in DMEM to inhibit protein synthesis or 0.0281% DMSO (control). Subsequently, cells were harvested at different time intervals post-CHX treatment (0, 8, 12, 24, 32, 48 h) for protein extraction, followed by western blot analysis. The bands corresponding to MTHFD2 were quantified utilizing ImageJ software, with actin bands employed for normalization. The half-life of MTHFD2 was determined by transforming the data to the natural logarithm (ln), performing a linear regression for each cell line, and dividing ln(0.5) by the slope of the corresponding regression equation.

#### Mouse Xenograft Model

All animal procedures complied with the institutional animal care use committee and relevant guidelines at Ben-Gurion University (permission number: 71-10-2023-E). Mice were housed in specific-pathogen-free (SPF) conditions at the Ben-Gurion University facility. All efforts were made to minimize animal suffering and ensure humane treatment.

A mouse xenograft model was employed to investigate the effects of HSPD1 KD on tumor growth. A total of 1 × $10^6$ A549 control cells or HSPD1KD cells, resuspended in 100 µL PBS, were injected subcutaneously into the left and right flanks of each 8-week-old female NOD.CB17-$Prkdc^{scid}$ mouse (*n* = 5 mice per group). At the end of the experiment, 37 days post-injection, mice were euthanized using $CO_2$ asphyxiation as IACUC guidelines. Tumors were excised and processed for downstream analysis. A portion of each tumor was snap-frozen in liquid nitrogen and then lysed to evaluate HSPD1 and MTHFD2 protein levels via western blot analysis. Another portion was fixed in 4% paraformaldehyde (PFA) and processed for proximity ligation assay (PLA) to assess molecular interactions using microscopy.

#### Mitochondrial mass measurements

Mitochondrial mass was measured using mitotracker™ green (FM- Invitrogen). Cells were plated in six-well plates (300,000 cells in each well) overnight; next, cells were washed and stained using mitotracker™ green. Next, cells were harvested, followed by centrifugation and resuspension of cells with 1 mll of PBS. Fluorescence was measured using a flow cytometer (Sysmex) and analyzed by FCS Express 5 (De Novo Software).

#### Mitochondrial membrane potential (MMP) measurements

Tetramethylrhodamine ethyl ester (TMRE) was used to measure mitochondria (MMP). Cells were plated in six-well plates (300,000 cells in each well) overnight. Next day, the mitochondrial uncopler Carbonyl cyanide-4-(trifluoromethoxy)phenylhydrazone (FCCP, 20mM) was added to the FCCP-control well for 10 min before adding TMRE (0.5µM). Cells were incubated for 20 minutes at 37°C. Next, cells were washed once with PBS and collected after ten minutes of trypsinization (200µl) and resuspension with (800µl) PBS. Cells were centrifuged once, and the pellet was resuspended with one ml of PBS. Fluorescence was measured using a flow cytometer (Sysmex) and analyzed by FCS Express 5 (De Novo Software).

#### SILAC

Cells were seeded at low density in 10cm dishes and allowed to grow for four passages. The shCONT cells were cultured in a medium containing light amino acids, while the HSPD1 KD cells were cultured with heavy amino acids. The light-medium contained L-Lysine-2HCl and L-Arginine-HCl, whereas the heavy medium included 13C6 L-Lysine-2HCl and 13C6

L-Arginine-HCl (0.1mg/ml). Upon reaching 90% confluency, the cells were divided into three 15cm plates and infected with lentiviruses targeting HSPD1 (shCONT, sh#1, and sh#2, respectively). For three additional passages, the cells were maintained under SILAC conditions (heavy or light amino acids). Approximately $2*10^7$ cells from each cell line were subsequently processed to extract mitochondrial proteins.

Mitochondrial isolation was carried out using MACS magnetic beads (Miltenyi Biotec) and slightly modified from manufacturer instructions. Following two washes with ice-cold PBS, cells were lysed with 1 mL of ice-cold lysis buffer per 10^7 cells in the dish. The lysis buffer consisted of 10 mL of 1x PBS and a Protease inhibitor cocktail tablet (1:10000). Cells were scraped off the dish, sonicated on ice for 15 seconds at 20% Amplitude, and then centrifuged at 1000g for 1 minute at 4°C. The supernatant was transferred to a 15 mL conical tube and mixed with 9 mL of ice-cold 1× PBS. Anti-TOM22 MicroBeads (50 µL) were added, and the mixture was incubated for 1 hour at 4 °C with gentle shaking. Subsequently, the lysate was applied stepwise onto LS Columns in a magnetic field, washed, and the magnetically labeled mitochondria were collected. The collected mitochondria were further processed by centrifugation, and the mitochondrial pellet was washed twice with PBS before final collection. The mitochondrial pellet was lysed using RIPA buffer, and the PierceTM BCA Protein Assay Kit was used to quantify the protein.

Soluble mitochondrial lysates were equally mixed. Control cell lysates, fed with light amino acids, were mixed with heavy isotope-labeled lysates from HSPD1 KD1 or KD2 (mix1 and mix2, respectively). Proteins were then identified and quantified using LCMS proteomics services provided by the Proteomic Unit at the Smoler Proteomic Center of the Technion.

The mass spectrometry proteomics data have been deposited to the ProteomeXchange Consortium via the PRIDE[1] partner repository with the dataset identifier PXD066753.

#### *In vitro* synthesis of isotopically labeled S-adenosyl-methionine$^{13}C_{10}$.

In the A549 experiment, SAM-$^{13}C_{10}$ was synthesized *in vitro* by Methionine adenosyltransferase (MAT,12µM) from methionine and Adenosine-$^{13}C_{10}$ 5′-triphosphate (ATP-$^{13}C_{10}$). The enzymatic reaction was performed for 2.5 hours at 37°C in a dry bath. The reaction was quenched using Perchloric acid (5%) followed by centrifugation at maximum speed at 4oC for 30 minutes. The supernatant was collected and transferred to HPLC tubs. 30µl of injection volume was loaded on the HPLC column (Chromatoggraphie-Service GMbH, #5861236). A total volume of 920µl was loaded. Each injection was analyzed for 18 minutes and read at 254nM. Fractions containing SAM-13C10 were collected manually and stored at -80°C. Samples were dried using a Lyophilizer. Subsequent samples were reconstituted and mixed with a total volume of 300µl sulfuric acid (10mM) to an estimated concentration of 3mM, based on the known concentrations of the substrates and stoichiometry in the *in vitro* enzymatic reaction. Calculating the exact concentration was performed in two steps. First, by o.d measurement at 260nm using nanodrop. Concentration was calculated using the molecule's molar absorption coefficient (ε=15,400/M*cm) and the Beer-Lambert law C=A/εl. A= absorbance at 260nm, l=1cm. This concentration is for the SAM-$^{13}C_{10}$ and its other molecules. This information is used to estimate purity levels. In the second step, HPLC measurement was performed along with a calibration curve of known concentrations of SAM. Samples were prepared with 1:1 perchloric acid (10%). The areas under the peak were calculated, and the concentration was calculated using the slope of the calibration curve. Samples were stored at -80ºC.

#### Metabolic extraction for liquid chromatography-mass spectrometry (LCMS)

Control and two KD cell lines were seeded in 6cm plates at 37 °C overnight (400,000 cells per well, minimum of six plates from each cell line). An additional 12-well plate with 100,000 cells per well (four wells for each cell line) was prepared for normalization by crystal violet staining. On the next day, metabolite extraction was performed as follows. First, the media was aspirated, and the cells were washed twice with 1ml of HPLC grade saline, ensuring complete saline removal before proceeding to the next step. Liquid nitrogen was poured onto the plate and placed in a liquid nitrogen tank. Plates were placed on ice for the extraction of metabolites. Metabolites were extracted with 1ml of a solution containing 50% methanol, 30% acetonitrile, and 20% water, all of which were of HPLC grade. Next, the plates were placed on an orbital shaker for 10 minutes at 4 °C. The extraction solution from each plate was then transferred into Eppendorf tubes and vortexed. Subsequently, the samples were centrifuged at 16,100g for 10 minutes at 4 °C. After centrifugation, all the supernatants were transferred to HPLC glass vials and were stored at -80 °C prior to LC-MS analysis.

#### SAM measurement

The dried samples were dissolved in 100uL of 80% methanol and 20% water (LCMS grade) using sonication for 5 minutes, ensuring complete dissolution. Subsequently, the solution was centrifuged for 10 minutes at maximum rpm, and the liquid was carefully transferred to new tubes. Following another centrifugation step for 10 minutes at maximum rpm, 80uL of the supernatant was carefully transferred to LC-MS vials, and the lids were securely closed. Reconstituted samples were analysed by LC-MS/MS. Ultra-performance liquid chromatography coupled with high-resolution mass spectrometry (UPLC-HRMS) analysis was conducted using Waters Acquity UPLC equipped with HSS-T3 column and connected to Thermo Q-Exactive HRMS equipped with electrospray ionization (ESI) source. The gradient of mobile phases A (0.1% formic acid in water) and B (acetonitrile) was used in the following order: 0-1 min 99.8% of A, 1-6 min 90% of A, 6-7 min 0% of A, 7-7.5 min 0% of A, 7.5-8 min 99.8% of A, 8-10 min 99.8% of A. The flow rate was 0.3 mL/min, temperature of the column was 40°C. The injection volume was 1µL. The MS parameters (spray voltage, vaporizer temperature, ion transfer tube, sheath gas) were set according to the manufacture recommendations. Data acquisition was performed in the full scan (70-700 m/z) positive ionization mode under the control of Xcalibur software, v. 4.5 (Thermo Scientific). The relative abundance of the SAM was calculated as a ration between peak area of the SAM and peak area of the D3-SAM internal standard.The area under the peak was calculated and normalized relative to crystal violet staining and heavy isotopic-labeled SAM.

#### Crystal violet staining

Cells were washed twice with PBS. The plates were washed and air-dried. Crystal Violet was added for half an hour, after which additional rinsing was performed. Following dehydration, 10% acetic acid was added to dissolve the Crystal Violet, and the optical density (OD) was measured at 590 nm.

#### Oxygen consumption rate (ORC)

ORC was measured by the Seahorse XF96 analyzer (Agilent). 6000 cells were seeded in 96-well XF cell culture plates with a final volume of 80 µL in each well (confluency of 60-90%). The seeded plates were then placed in the incubator overnight. Next day: the wells were washed, and the medium was replaced by an OCR assay medium supplemented with glucose (10 mM), sodium pyruvate (1 mM), and glutamine (2 mM). Washing and replacing the medium with OCR assay medium as follows: 60 µL of the medium was removed from each well, followed by adding 200 µL of assay medium to the well. Subsequently, 200 µL of the medium was removed from the well again, and 160 µL of assay medium was added, resulting in a final volume of 180 µL in each well. The cells were then incubated for 45-60 minutes in the incubator. Next, the ports were injected with the proper inhibitor according to manufacturer instructions. Oligomycin (1.5µM), FCCP(1µM), Rot/AA (0.5µM), and OCR assay medium(28µl) were injected into ports A-D respectively. After calibration, the assay plate was placed in the Seahorse XF96 analyzer for OCR measurements. Basal and maximal respiration rates were calculated by averaging the normalized OCR values of six replicates at three different time points. The experiment was performed twice. 4′,6-diamidino-2-phenylindole (DAPI) DNA staining was used to normalize the OCR reads to the relative cell count.

#### The purification of MTHFD2

The cDNA of the MTHFD2 sequence was purchased from Sino Biological. The sequence included the mitochondrial leader peptide (AA1-35)153. Using PCR amplification, the cDNA of MTHFD2 lacking its mitochondrial lea leader peptide (AA36-350) _6*Histag was clones to pET-24, a cloning vector, and expressed in E.coli (Bl21_DE3). Isopropyl β-d-1-thiogalactopyranoside (IPTG, 0.1mM) for 16 hours at 30 °C was used to induce vector expression for three hours at 30 °C. Next, call lysates were loaded onto a nickel column (Hits Trap HP column, 5ml, GE Healthcare), followed by washing with wash buffer and eluted with elution buffer. Fractions were collected and loaded on 15% SDS-PAGE for gel electrophoresis. Fractions containing the MTHFD2 were dialyzed and subjected to size exclusion chromatography. Fractions containing MTHFD2 were concentrated using Amicon (Millipore, ACS501024). BCA was used to determine the concentration of purified MTHFD2. MTHFD2 was aliquoted and stored at -80oC.

#### MTHFD2 activity assay

MTHFD2 activity was assessed as previously described with some modifications[146,154].180µl of enzyme buffer containing 20nM or 10nM of MTHFD2 was loaded into a clear 96-well plate (Corning, CLS9018BC-100EA) and placed inside a plate reader. Next, 20ul of activity buffer containing different concentrations of the substrate 5,10-Methenyl-tetrahydrofolate (Schircks Laboratories) was added simultaneously to the plate using a multi-channel pipettor (1:10 ratio). Reads at 350nm and 30oC were taken immediately after in five-second intervals for ten minutes. For refolding by chaperone assay, 20nM of enzyme and 0.4mM of substrate were used.

#### Chaperone-assisted protein folding assay

The MTHFD2 refolding assay was performed as described previously, with some modifications. MTHFD2 assay MTHFD2 was denatured using 30mM of HCl for one hour. Next, denatured MTHFD2 was transferred to the binding buffer, containing with or without the chaperone (HSPD1/GroEL) for 30 minutes 30°C (1:25 dilution ratio). Next, the MTHFD2-chaperone complex was transferred to a folding buffer with or without ATP and co-chaperone (HSPE1/GroES, 1:1 dilution ratio). The ratio of 2:14:14 (MTHFD2:chaperone:co-chaperone) was determined based on previously published stoichiometries[2,3]. After one hour of incubation at room temperature, the activity of MTHFD2 was assessed as described above. The rate of accumulation of the Methenyl-THF was analyzed by calculating the slope of the curve representing optical density (O.D.) at 350 nm as a function of time. This analysis was performed using KaleidaGraph (Synergy Software). The slopes were used as an indicator for MTHFD2 activity. The activity of native MTHFD2 was used to determine 100% activity. HSPD1 and HSPE1 were a kind gift from Prof. Abdussalam Azem of Tel-Aviv University.

#### RNA sequencing

Total RNA was extracted from cells using an RNA purification kit (Invitrogen), following the manufacturer's instructions. RNA concentration and quality were determined using a NanoDrop spectrophotometer. Samples were then sent to the sequencing unit in Ben-Gurion for library preparation and sequencing. RNA integrity was evaluated using a QIAxcel device (QIAGEN, QIAxcel RNA QC Kit v2.0), and RNA concentration was measured using the QuantiFluor® RNA System (Promega, #E3310) on a Qubit™ Flex Fluorometer (Invitrogen). From 0.5 µg of total RNA, mRNA was enriched using the NEBNext® Poly(A) mRNA Magnetic Isolation Module (New England Biolabs, #E7490), ensuring specific capture of polyadenylated transcripts. According to the manufacturer's instructions, stranded RNA-seq libraries were prepared using the NEBNext® Ultra™ II Directional RNA Library Prep Kit for Illumina® (New England Biolabs, #E7760). The molarity of RNA-seq libraries was determined using the QIAxcel device (QIAxcel DNA High Sensitivity Kit) and the QuantiFluor® dsDNA System (Promega, #E2670) on a Qubit™ Flex Fluorometer (Invitrogen). Libraries were sequenced on a NovaSeq X system (Illumina) with paired-end reads of 150 base pairs (150PE). Sequencing data were processed by Linor Cohen using a bioinformatics pipeline implemented in the software NeatSeq-Flow[4].

##### Differential Expression Analysis

Gene-level differential expression analysis was performed using the NeatSeq-Flow platform DESeq2[5] module. Expression matrices from RSEM were used as input. The statistical model included a batch as well as genotype and fraction interaction, predefined contrasts tested fraction comparisons within each cell line separately. Genes with fold change ≥1, adjusted P-value <.05 were considered as significantly differentially expressed genes. Significant genes were clustered using the 'eclust' function from the factorextra R package[6]. Enrichment for Gene Ontology biological processes and KEGG pathways was performed using clusterProfiler v4.0 R package[7].

Sequencing data were processed by Linor Cohen using a bioinformatics pipeline implemented in the software NeatSeq-Flow.

#### *C. elegans* strains and maintenance

All *C. elegans* strains used in this work are listed in Table 7. Nematodes were maintained on NGM plates seeded with OP50-1 *Escherichia coli* bacteria at 15 °C. For each experiment, eggs were picked from plates maintained at 15 °C and transferred to 20 °C for the duration of the experiments.

To induce male formation, synchronized young adult hermaphrodites were placed on fresh NGM plates (5–10 worms per plate) and incubated at 32°C for 4 hours. Following this heat shock, plates were maintained at 20 °C for 3 days to allow recovery and development. Plates were then examined under a dissecting microscope for the presence of males. Male worms were subsequently used for crosses to generate experimental lines. The genotypes of crossed progeny were determined by PCR followed by gel electrophoresis and sequencing.

##### Live imaging preparation

For live imaging, worms were mounted on 2% agarose pads prepared on glass slides containing 1.5 mM levamisole for immobilization. Approximately 40 µl of agarose was placed on a glass slide and immediately covered with a coverslip to form the pad. Worms were transferred into a 1–2 µl drop of levamisole on the pad, covered with a coverslip, and imaged immediately after preparation to ensure accuracy of cellular structures.

##### Thrashing Assay (Liquid Motility)

Motility of age-synchronized worms was analyzed. An individual worm was transferred into 100 µL of M9 buffer in a plate and allowed to acclimate for 5 min. Motility was quantified by counting the number of body bends over a 20s interval under a stereomicroscope, and values were converted to body bends per minute. For comparison, CRISPR-generated mutant worms targeting *hsp-60* and *hsp-10* were analyzed, with SJ4100 UPRmt reporter worms used as the control strain.

#### Table 1. Materials

| Material | Source | Identifier |
|---|---|---|
| DMEM, high glucose | Gibco | Cat#11965092 |
| Fetal bovine serum (FBS) | ThermoFisher Scientific | Cat#10270106 |
| sodium pyruvate solution | Biological Industries | Cat#03-042-1 |
| antibiotic–antimycotic 100X | TOKU-E | Cat#A045 |
| Gene Knockout Kit v2 | SYNTHEGO | Cat#111 |
| Lipofectamine CRISPRMAX Cas9 transfection reagents | ThermoFisher Scientific | Cat#CMAX00001 |
| Opti-MEM I reduced serum medium. | ThermoFisher Scientific | Cat#31985062 |
| cDNA Synthesis Kit | New England BioLabs | Cat#E6300S |

| | | |
|---|---|---|
| LipoJet™ In Vitro Transfection Kit | SignaGen Laboratories | Cat#SL100468 |
| DAPI Fluoromount-G® | Southern Biotech | Cat#0100-20 |
| Dimethyl sulfoxide (DMSO) | Sigma-Aldrich | Cat#D8418 |
| Ponceau S solution | Sigma-Aldrich | Cat#P7170-1L |
| CalFectin™Mammalian Cell Transfection Reagent | SignaGen Laboratories | Cat#SL100478 |
| Protease Inhibitor Cocktail | Sigma-Aldrich | Cat#P8340 |
| PhosSTOP™ | Roche | Cat#4906837001 |
| Crystal violet | Sigma-Aldrich | Cat#C0775 |
| Cycloheximide (CHX) | Sigma-Aldrich | Cat#01810 |
| PBS | Invitrogen | Cat# 10010-023 |
| PFA | ENCO | Cat# 0219998320 |
| Triton X-100 | Sigma-Aldrich | Cat# 000010 |

| | | |
|---|---|---|
| Duolink® In Situ PLA® Probe Anti-Mouse MINUS | Sigma-Aldrich | Cat#DUO92004 |
| Duolink® In Situ PLA® Probe Anti-Rabbit PLUS | Sigma-Aldrich | Cat#DUO92002 |
| Duolink® In Situ Wash Buffers, Fluorescence | Sigma-Aldrich | Cat#DUO82049 |
| Duolink® In Situ Detection Reagents FarRed | Sigma-Aldrich | Cat#DUO92013 |
| Duolink® In Situ Detection Reagents FarRed | Sigma-Aldrich | Cat#DUO92013 |
| CalfectinTM Mammalian Cell Transfection Reagent | SignaGen | Cat#SL100478 |
| MitoTracker™ Red CMXRos | ThermoFisher Scientific | Cat#M7512 |
| MitoTracker™ Green FM | ThermoFisher Scientific | Cat#M7514 |
| qPCRBIO SyGreen® Blue Mix | PCR Biosystems | Cat#PB20.16-05 |
| Protease Inhibitor Cocktail | Sigma-Aldrich | Cat#P8340 |
| Methanol Analyzed LC-MS | J.T.Baker® | Cat#9822 |
| HPLC grade wated | Sigma-Aldrich | Cat#34877 |

| | | |
|---|---|---|
| SILAC Protein Quantification Kit | ThermoFisher Scientific | A33969 |
| 13C6 L-Arginine-HCl | ThermoFisher Scientific | 88210 |
| Sodium Formate | Sigma-Aldrich | 107603-1KG |
| Adenosine triphosphate $^{13}C_{10}$ 5 (ATP-$^{13}C_{10}$). | Sigma-Aldrich | 710695 |
| S-Adenosyl-L-methionine-d3 5mg | biotag | HY-B0617S |
| Amicon Ultra 0.5 ml Centrifugal filters | Sigma-Aldrich | UFC5003 |
| MACS, mitochondrial isolation KIT | Miltenyi Biotec | 130-094-532 |
| 5,10-Methylene-tetrahydrofolate | Schircks Laboratories | 16.226 |
| Agarose | Sigma-Aldrich | |
| Tetramisole hydrochloride | Sigma-Aldrich | L9756-5G |

#### Table 2. Antibodies

| Antibody | Source | Identifier |
|---|---|---|
| MTHFD2 (WB: 1:1000) | Abcam | Cat#ab151447 |

| | | |
|---|---|---|
| HSPD1 (WB: 1:1000) | Santa Cruz | Cat#sc-13115 |
| Citrate synthase (WB: 1:1000) | Abcam | Cat# ab96600 |
| HSPA9 (WB: 1:1000) | Cell Signaling Technology | Cat#sc-133137 |
| Anti-mouse IgG (WB: 1:1500) | Cell Signaling Technology | Cat#7076 |
| Anti-rabbit IgG (WB: 1:1500) | Cell Signaling Technology | Cat#7074 |
| Actin (WB: 1:10000) | Merck | Cat# MAB1501 |
| HSPE1 (WB: 1:1000) | Santa Cruz | Cat#sc-376313 |
| HSC-70 (WB: 1:5000) | Santa Cruz | Cat#sc-7298 |

#### Table 3. shRNA and siRNA sequences

| | |
|---|---|
| shControl (for HSPD1 KD) | CCTAAGGTTAAGTCGCCCTCGCTCGAGCGAGGGCGACTTAACCTTAGG |
| shHSPD1 #1 | CCGGCCTGCTCTTGAAATTGCCAATCTCGAGATTGGCAATTTCAAGAGCAGGTTTTT |
| shHSPD1 #2 | CCGGGCAATGACCATTGCTAAGAATCTCGAGATTCTTAGCAATGGTCATTGCTTTTT |

| | |
|---|---|
| shControl (for HSPE1 KD) | ATCTCGCTTGGGCGAGAGTAAG |
| shHSPE1 #1 | AATCTTGGGCTAGGTTGTC |
| shHSPE1 #2 | AATAGAGGTTGAAAGTGCG |
| siControl (pool) | ACCAAAUGUACAGCUGAUU, ACCAAAUGUACAACACACU, ACCAAAUGUACAAAAGACU, ACCAAAUGUACAAAAGGAU |
| siLONP1 (pool) | CCUUUAGCCGGAUGUUCUA, CGCGUACAUACUGCACUGU, GCUCUCAUGAAGGCGAAGU, CUUCUCUGGUUAUAAGGAU |
| siClpP (pool) | CUCUCCCUGGCGAUGUACU, UUUGUCUCGGACGUCCACU, GUUGUCGGGACCACCACAC, UCUUCGGGUAGGUGUACAU |

##### Table 4. Plasmids

| Plasmid | Source | Identifier |
|---|---|---|
| pMD2.G | Addgene | Cat#12259 |
| psPAX2 | Addgene | Cat#12260 |
| MTHFD2 (AA36-350_histag) amino acid sequence.<br><br>Sequence. The first methionine is in blue—the stop codon is in red. The his-tag sequence is in bold. | **M**EAVVISGRK LAQQIKQEVR QEVEEWVAS GNKRPHLSVI LVGENPASHS YVLNKTRAAA VVGINSETIM KPASISEEELL NLINKLNNDD NVDGLLVQLP LPEHIDERRI CNAVSPDKD VDGFHVINVG RMCLDQYSM LPATPWGVW EIIKRTGIPTL GKNVVVAGR SKNVGMPIA MLLHTDGAH ERPGGDATV TISHRYTPKE QLKKHTILADI VISAAGIPNLI TADMIKEGAA VIDVGINRVH DPVTAKPKLV GDVDFEGVR QKAGYITPVP GGVGPMTVA MLMKNTIIAA KKVLRLEERE VLKSKELGVA TN**HHHHHH*** | |

#### Table 5. qRT-PCR primers

| gene | primer f | primer r |
|---|---|---|
| HSPD1 | GGTCTTCAGGTTGTGGCAGT | TTCAGGGTCAATCCCTCTTC |
| MTHFD2 | TGGGGTGTGTGGGAAATAAT | TGGGCATTCCAACGTTTT |
| LONP1 | ATCGTGAAGACCATCCGGGAC | ACAGCCGCTTAGGAATATTGG TCT C |
| ClpP | CCCGTATCATGATCCACCA | AGAGCTGCTTCTTGAGCTTCAT |
| L32 | GCACACTGACTACAGCCTTGA | TACCCAGGTTTGGAGGTGTG |

**Table 6. Buffers**

| Buffer | Component and final concentrations |
|---|---|
| RIPA lysis buffer | 150 mM NaCl; 50 mM Tris pH = 8.0; 1% Triton X-100; 0.5% sodium deoxycholate; 0.1% SDS |
| TBS | 20 mM Tris HCl, pH 7.4, 150 mM NaCl |
| Running buffer | 25 mM Tris base, 192 mM Glycine, 0.1% SDS, pH=8.3 |
| Transfer Buffer | 12 mM Tris base, 196mM Glycine, pH=8.3 |
| HPLC grade saline | 0.9% NaCl in HPLC grade $H_2O$ |

| | |
|---|---|
| X5 Sample buffer | 250 mM Tris, 40% Glycerol, 10% SDS, 0.25% bromophenol blue, pH 6.8 |
| SAM-$^{13}C_{10}$ synthesis | 25mM Tris-Hcl, p.H 8, 100mM KCl, 10mM mgCl2, 1mM DDT, L-Methionine 2mM, ATP-$^{13}C_{10.}$ Methionine adenosyltransferase (12μM) |
| HPLC running buffer | 400mM Amonium formate, 0.53% formic acid pH 4 |
| HPLC grade saline | 0.9% NaCl in HPLC grade $H_2O$ |
| Equilibration Buffer | 50mM Tris-Hcl, p.H 8.5, 300mM NaCl, 10% glycerol, 10mM imidazole, 5mM β-mercaptoethanol. |
| Wash Buffer | 50mM Tris-Hcl, p.H 8.5, 300mM NaCl, 10% glycerol, 20mM imidazole, 5mM β-mercaptoethanol. |
| elution buffer | 50mM Tris-Hcl, p.H 8.5, 300mM NaCl, 10% glycerol, 250mM imidazole, 5mM β-mercaptoethanol. |
| Size exclusion buffer | 50mM Tris-Hcl, p.H 8.5, 300mM NaCl |
| MTHFD2 storgae buffer | 50mM Tris-Hcl, p.H 8.5, 300mM NaCl |
| MTHFD2 enzyme buffer | 50mM HEPES ,pH 8, ,25 mM potassium phosphate, pH 7.3, |

| | |
|---|---|
| | 100mM KCl, 5 mM MgCl2, 40mM β-mercaptoethanol<br>1mM NAD |
| MTHFD2 activity buffer | 50mM HEPES ,pH 8, ,25 mM potassium phosphate, pH 7.3, 100mM KCl, 5 mM MgCl2, 40mM β-mercaptoethanol<br>1mM NAD, Metylene-THF 0.4mM |
| MTHFD2-HSPD1 binding buffer | 65mM potassium phosphate, 20mM MgCl2, 50mM KCl<br>5mM DTT, *1.2μM HSPD1(*when presents) |
| Folding co-HSPE1 buffer | 65mM potassium phosphate, 20mM MgCl2, 50mM KCl,<br>5mM DTT, *2mM ATP,*20 units/mL PK, *20mM PEP,<br>*0.8μM HSPE1 (*when presents) |
| MTHFD2-GroEs binding buffer | 50mM triethanolamine, 20mM MgCl2, 150mM KCl, 1mM DTT, 10 units/mL PK, 10mM PEP, 1mM ATP, 1.2μM groEL |
| Folding-GroEl binding buffer | 50mM triethanolamine, 20mM MgCl2, 150mM KCl, 1mM DTT, 10 units/mL PK, 10mM PEP, 1mM ATP, 0.8μM groEL |
| M9 buffer | 22 mM KH2PO4, 42 mM Na2HPO4, 85 mM NaCl, 1 mM MgSO4 |

#### Table 7. Experimental models Organism/Strain

| Organism/Strain | Source | Identifier |
|---|---|---|
| Mouse: NOD *SCID* Gamma *Prkdc*$^{scid}$ | The Jackson Laboratory | RRID: IMSR_JAX:001303 |
| *C. elegans:* N2, SJ4100, VH7154, VC4728 | CGC, which is funded by NIH Office of Research Infrastructure Programs (P40 OD010440) | |

#### Original blots

Figure 1A:

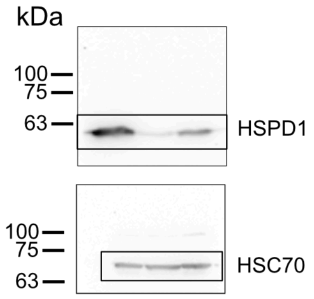


These original blots belongs to Figure 1F

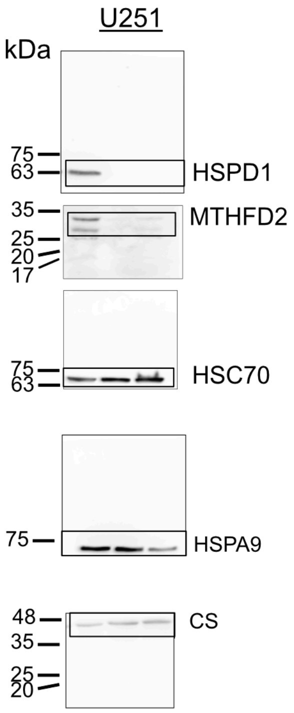


These original blots belongs to Figure 1G

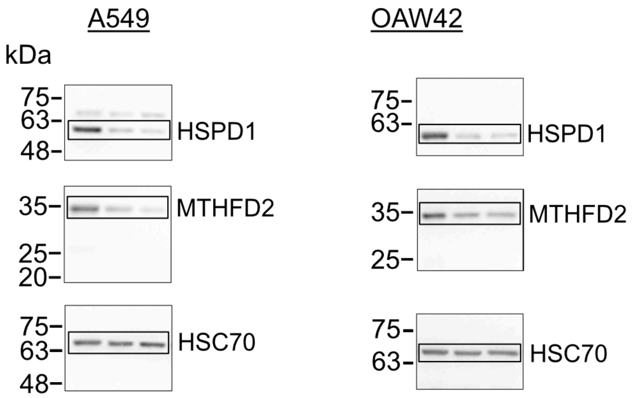

These original blots belongs to Figure 3C

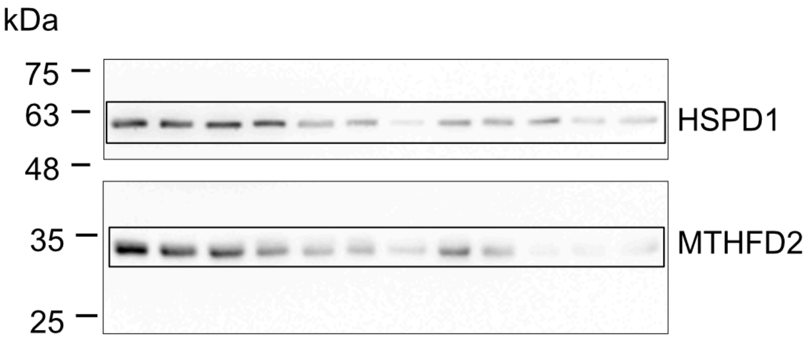


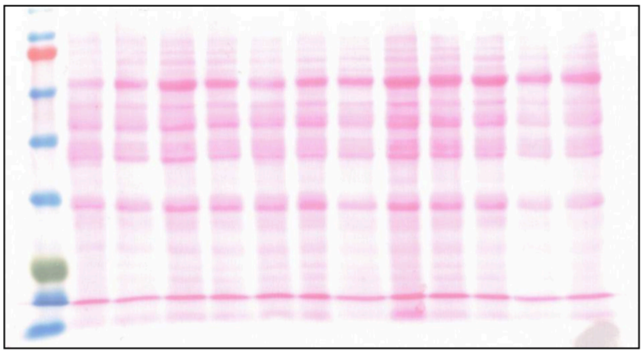

These original blots belongs to Figure 4A

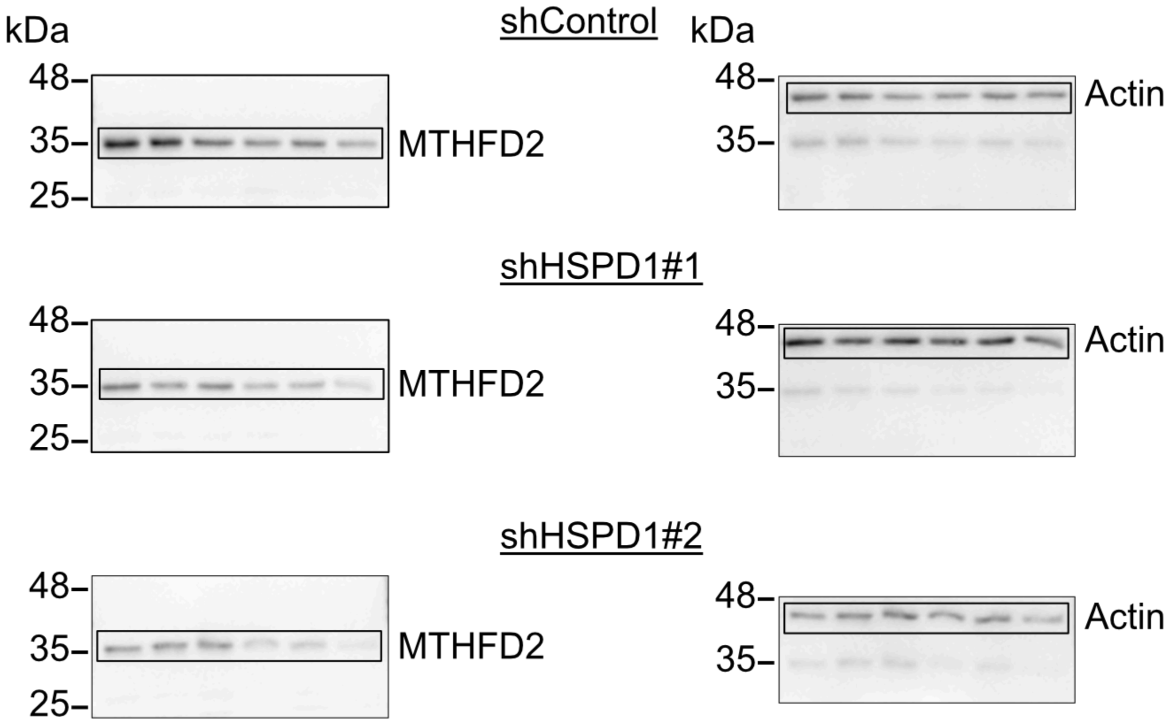


These original blots belongs to Figure 4C

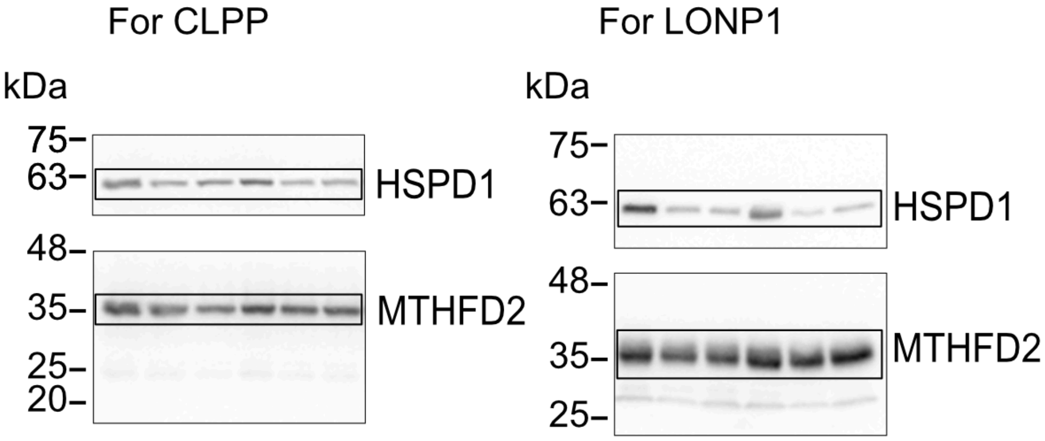

These original blots belongs to Figure 5C

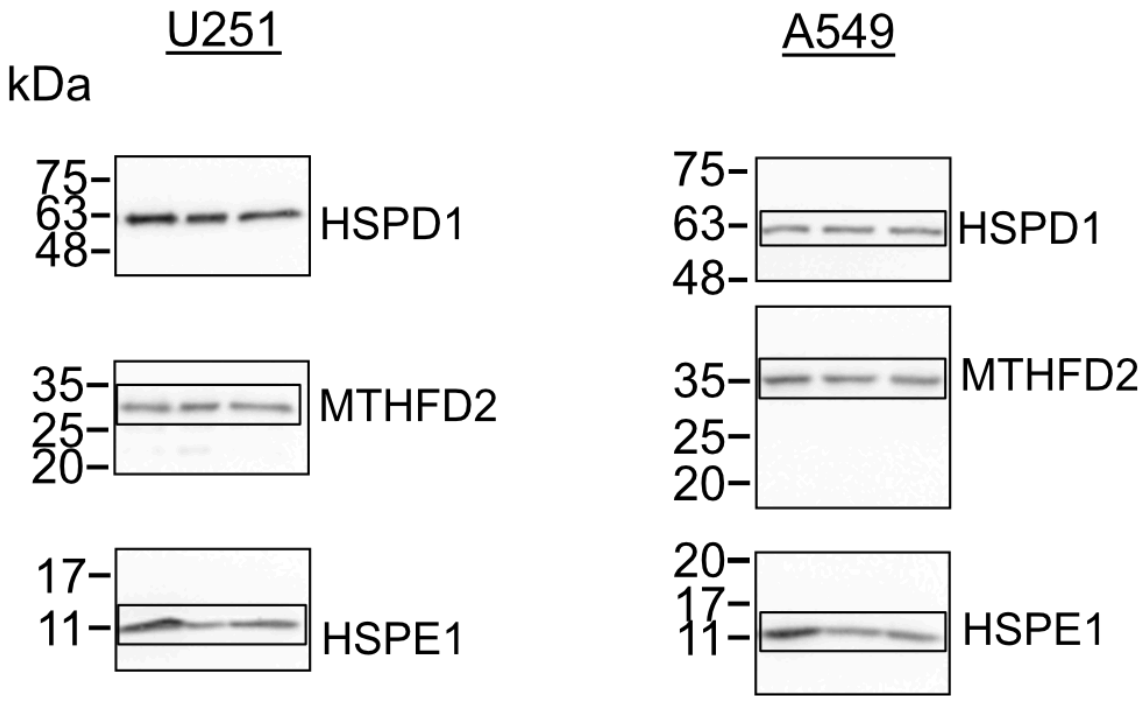
